# GET_PHYLOMARKERS, a software package to select optimal orthologous clusters for phylogenomics and inferring pan-genome phylogenies, used for a critical geno-taxonomic revision of the genus *Stenotrophomonas*

**DOI:** 10.1101/294660

**Authors:** Pablo Vinuesa, Luz Edith Ochoa-Sánchez, Bruno Contreras-Moreira

## Abstract

The massive accumulation of genome-sequences in public databases promoted the proliferation of genome-level phylogenetic analyses in many areas of biological research. However, due to diverse evolutionary and genetic processes, many loci have undesirable properties for phylogenetic reconstruction. These, if undetected, can result in erroneous or biased estimates, particularly when estimating species trees from concatenated datasets. To deal with these problems, we developed GET_PHYLOMARKERS, a pipeline designed to identify high-quality markers to estimate robust genome phylogenies from the orthologous clusters, or the pan-genome matrix (PGM), computed by GET_HOMOLOGUES. In the first context, a set of sequential filters are applied to exclude recombinant alignments and those producing anomalous or poorly resolved trees. Multiple sequence alignments and maximum likelihood (ML) phylogenies are computed in parallel on multi-core computers. A ML species tree is estimated from the concatenated set of top-ranking alignments at the DNA or protein levels, using either FastTree or IQ-TREE (IQT). The latter is used by default due to its superior performance revealed in an extensive benchmark analysis. In addition, parsimony and ML phylogenies can be estimated from the PGM.

We demonstrate the practical utility of the software by analyzing 170 *Stenotrophomonas* genome sequences available in RefSeq and 10 new complete genomes of environmental *S. maltophilia* complex (Smc) isolates reported herein. A combination of core-genome and PGM analyses was used to revise the molecular systematics of the genus. An unsupervised learning approach that uses a goodness of clustering statistic identified 20 groups within the Smc at a core-genome average nucleotide identity of 95.9% that are perfectly consistent with strongly supported clades on the core- and pan-genome trees. In addition, we identified 14 misclassified RefSeq genome sequences, 12 of them labeled as *S. maltophilia*, demonstrating the broad utility of the software for phylogenomics and geno-taxonomic studies. The code, a detailed manual and tutorials are freely available for Linux/UNIX servers under the GNU GPLv3 license at https://github.com/vinuesa/get_phylomarkers. A docker image bundling GET_PHYLOMARKERS with GET_HOMOLOGUES is available at https://hub.docker.com/r/csicunam/get_homologues/, which can be easily run on any platform.

## INTRODUCTION

Accurate phylogenies represent key models of descent in modern biological research. They are applied to the study of a broad spectrum of evolutionary topics, ranging from the analysis of populations up to the ecology of communities (Dornburg et al., 2017). The way microbiologists describe and delimit species is undergoing a major revision in the light of genomics (Rosselló-Móra and Amann, 2015; Vandamme and Peeters, 2014), as reflected in the emerging field of microbial genomic taxonomy (Konstantinidis and Tiedje, 2007; Thompson et al., 2009, 2013). Current geno-taxonomic practice is largely based on the estimation of (core-)genome phylogenies (Ciccarelli et al., 2006; Daubin et al., 2002; Lerat et al., 2003; Tettelin et al., 2005; Wu and Eisen, 2008) and the computation of diverse overall genome relatedness indices (OGRIs) (Chun and Rainey, 2014), such as the popular genomic average nucleotide identity (gANI) values (Goris et al., 2007; Konstantinidis and Tiedje, 2005; Richter and Rossello-Mora, 2009). These indices are rapidly and effectively replacing the traditional DNA-DNA hybridization values used for species delimitation in the pre-genomic era (Stackebrandt et al., 2002; Stackebrandt and Goebel, 1994; Vandamme et al., 1996).

The ever-increasing volume of genome sequences accumulating in public sequence repositories provides a huge volume of data for phylogenetic analysis. This significantly improves our capacity to understand the evolution of species and any associated traits (Dornburg et al., 2017). However, due to diverse evolutionary forces and processes, many loci in genomes have undesirable properties for phylogenetic reconstruction. If undetected, these can lead to erroneous or biased estimates (Parks et al., 2018; Shen et al., 2017), although, ironically, with strong branch support (Kumar et al., 2012). Their impact is particularly strong in concatenated datasets (Degnan and Rosenberg, 2009; Kubatko and Degnan, 2007), which are standard in microbial phylogenomics (Wu and Eisen, 2008). Hence, robust phylogenomic inference requires the selection of well-suited markers for the task (Vinuesa, 2010).

For this study we developed GET_PHYLOMARKERS, an open-source and easy-to-use software package designed with the aim of inferring robust genome-level phylogenies and providing tools for microbial genome taxonomy. We describe the implementation details of the pipeline and how it integrates with GET_HOMOLOGUES (Contreras-Moreira and Vinuesa, 2013; Vinuesa and Contreras-Moreira, 2015). The latter is a popular and versatile genome-analysis software package designed to identify robust clusters of homologous sequences. It has been widely used in microbial pan-genomics and comparative genomics (Lira et al., 2017; Nourdin-Galindo et al., 2017; Sandner-Miranda et al., 2018; Savory et al., 2017), including recent bacterial geno-taxonomic (Gauthier et al., 2017; Gomila et al., 2017), and plant pan-genomic studies (Contreras-Moreira et al., 2017; Gordon et al., 2017). Regularly updated auxiliary scripts bundled in the GET_HOMOLOGUES package compute diverse OGRIs, at the protein, CDS and transcript levels, provide graphical and statistical tools for a range of pan-genome analyses, including inference of pan-genome phylogenies under the parsimony criterion. GET_PHYLOMARKERS was designed to work both at the core-genome and pan-genome levels, using either the homologous gene clusters or the pan-genome matrix computed by GET_HOMOLOGUES. In the first context, it identifies single-copy orthologous gene families with optimal attributes (listed further down) and concatenates them to estimate a genomic species tree. In the second scenario, it uses the pan-genome matrix (PGM) to estimate phylogenies under the maximum likelihood (ML) and parsimony optimality criteria. In addition, we implemented unsupervised learning methods that automatically identify species-like genome clusters based on the statistical analysis of the PGM and core-genome average nucleotide identity matrices (cgANIb).

To demonstrate these capabilities and benchmark performance, we applied the pipeline to critically evaluate the molecular systematics and taxonomy of the genus *Stenotrophomonas.* Species delimitation is problematic and far from resolved in this genus (Ochoa-Sánchez and Vinuesa, 2017), despite recent efforts using genomic approaches with a limited number of genome sequences (Lira et al., 2017; Patil et al., 2016; Yu et al., 2016).

The genus *Stenotrophomonas* (Gammaproteobacteria, *Xhanthomonadales*, *Xanthomonadaceae*) (Palleroni, 2005; Palleroni and Bradbury, 1993) groups ubiquitous, aerobic, non-fermenting bacteria that thrive in diverse aquatic and edaphic habitats, including human-impacted ecosystems (Ryan et al., 2009). As of March 2018, 14 validly described species were listed in Jean Euzeby’s list of prokaryotic names with standing in nomenclature (http://www.bacterio.net/stenotrophomonas.html). By far, its best-known species is *S. maltophilia.* It is considered a globally emerging, multidrug-resistant (MDR) and opportunistic pathogen (Brooke, 2012; Chang et al., 2015). *S. maltophilia*-like organisms display high genetic, ecological and phenotypic diversity (Valdezate et al., 2004; Vasileuskaya-Schulz et al., 2011), forming the so-called *S. maltophilia* complex (Smc) (Berg and Martinez, 2015; Svensson-Stadler et al., 2012). Heterogeneous resistance and virulence phenotypes have been reported for environmental isolates of diverse ecological origin classified as *S. maltophilia* (Adamek et al., 2011; Deredjian et al., 2016). We have recently shown that this phenotypic heterogeneity largely results from problems in species delimitations within the Smc (Ochoa-Sánchez and Vinuesa, 2017). We analyzed the genetic diversity in a collection of 108 *Stenotrophomonas* isolates recovered from several water bodies in Morelos, Central Mexico, based on sequence data generated for the 7 loci used in the Multilocus Sequence Typing (MLST) scheme available for *S. maltophilia* at https://pubmlst.org. We assembled a large set of reference sequences retrieved from the MLST database (Kaiser et al., 2009; Vasileuskaya-Schulz et al., 2011) and from selected genome sequences (Crossman et al., 2008; Davenport et al., 2014; Lira et al., 2012; Patil et al., 2016; Vinuesa and Ochoa-Sánchez, 2015), encompassing 11 out of the 12 validly described species at the time. State-of-the-art phylogenetic and population genetics methods, including the multispecies coalescent model coupled with Bayes factor analysis and Bayesian clustering of the multilocus genotypes consistently resolved five conservatively-defined genospecies within the Smc clade, which were named *S. maltophilia* and Smc1-Smc4. The approach also delimited Smc5 as a sister clade of *S. rhizophila*. Importantly, we showed that *i*) only members of the Smc clade that we designed as *S. maltophilia* were truly MDR and *ii*) that *S. maltophilia* was the only species that consistently expressed metallo-beta-lactamases (Ochoa-Sánchez and Vinuesa, 2017). Strains of the genospecies Smc1 and Smc2 were only recovered from the Mexican rivers and displayed significantly lower resistance levels than sympatric *S. maltophilia* isolates, revealing well-defined species-specific phenotypes.

Given this context, the present study was designed with two major goals. The first one was to develop GET_PHYLOMARKERS, a pipeline for the automatic and robust estimation of genome phylogenies using state-of-the art methods. The emphasis of the pipeline is on selecting top-ranking markers for the task, based on the following quantitative/statistical criteria: *i*) they should not present signs of recombination, *ii*) the resulting gene trees should not be anomalous or deviating from the distribution of tree topologies and branch lengths expected under the multispecies coalescent model and *iii*) they should have a strong phylogenetic signal. The top-scoring markers are concatenated to estimate the species phylogeny under the maximum likelihood optimality criterion using either FastTree (Price et al., 2010) or IQ-TREE (Nguyen et al., 2015). The second aim was to apply GET_PHYLOMARKERS to challenge and refine the species delimitations reported in our previous MLSA study (Ochoa-Sánchez and Vinuesa, 2017) using a genomic approach, focusing on resolving the geno-taxonomic structure of the Smc and *S. maltophilia sensu lato* clades. For this purpose we sequenced five strains from the new genospecies Smc1 and Smc2 and analyzed them together with all reference genome sequences available for the genus *Stenotrophomonas* as of August 2017 using the methods implemented in GET_PHYLOMARKERS. The results were used to critically revise the molecular systematics of the genus in light of genomics, identify misclassified genome sequences, suggest correct classifications for them and discover multiple novel genospecies within *S. maltophilia*.

## MATERIALS AND METHODS

### Genome sequencing, assembly and annotation

Ten *Stenotrophomonas* strains from our collection were selected (Table 1) for genome sequencing using a MiSeq instrument (2x300 bp) at the Genomics Core Sequencing Service provided by Arizona State University (DNASU). They were all isolated from rivers in the state of Morelos, Central Mexico, and classified as genospecies 1 (Smc1) or 2 (Smc2), as detailed in a previous publication (Ochoa-Sánchez and Vinuesa, 2017). Adaptors at the 5’-ends and low quality residues at the 3’ ends of reads were trimmed-off using ngsShoRT v2.1 (Chen et al., 2014) and passed to Spades v3.10.1 (Bankevich et al., 2012) for assembly (with options --careful -k 33,55,77,99,127,151). The resulting assembly scaffolds were filtered to remove those with low coverage (< 7X) and short length (< 500 nt). All complete genome sequences available in RefSeq for *Stenotrophomonas* spp. were used as references for automated ordering of assembly scaffolds using MeDuSa v1.6 (Bosi et al., 2015). A final assembly polishing step was performed by remapping the quality-filtered sequence reads on the ordered scaffolds using BWA (Li and Durbin, 2009) and passing the resulting sorted binary alignments to SAMtools (Li et al., 2009) for indexing. The indexed alignments were used by Pilon 1.21 (Walker et al., 2014) for gap closure and filling, correction of indels and single nucleotide polymorphisms (SNPs), as previously described (Vinuesa and Ochoa-Sánchez, 2015). The polished assemblies were annotated with NCBI’s Prokaryotic Genome Annotation Pipeline (PGAP v4.2) (Angiuoli et al., 2008). BioProject and BioSample accession numbers are provided in Table S1.

**Table 1.** Overview of key annotation features for the 10 new genome assemblies reported in this study for environmental isolates recovered from Mexican rivers and classified as genospecies 1 (Smc1) and 2 (Smc2) in the study of Ochoa-Sánchez and Vinuesa (2017). Details of their isolation sites and antimicrobial resistance phenotypes can be found therein. All genomes consist of a single gapped chromosome. Supplementary table S1 provides additional information of the assemblies. Their phylogenetic placement within the *Stenotrophomonas maltophilia* complex is shown in Figure 5 (clades Sgn1/Smc1 and Sgn2/Smc2).

| Genome | Size_nt | CDSs<br>(coding) | rRNAs | tRNAs | pseudo-<br>genes | RefSeq<br>Acc. num. |
| --- | --- | --- | --- | --- | --- | --- |
| *Stenotrophomonas* genospecies 1<br>(Smc1; Sgn1) ESTM1D MKCIP4 1 | 4,475,880 | 3,904 | 6 | 59 | 67 | CP026004 |
| *Stenotrophomonas* genospecies 1<br>(Smc1; Sgn1) SAU14A NAIMI4 5 | 4,570,883 | 4,020 | 6 | 69 | 66 | CP026003 |
| *Stenotrophomonas* genospecies 1<br>(Smc1; Sgn1) ZAC14A NAIMI4 1 | 4,698,328 | 4,150 | 7 | 45 | 66 | CP026002 |
| *Stenotrophomonas* genospecies 1<br>(Smc1; Sgn1) ZAC14D1 NAIMI4 1 | 4,702,461 | 4,131 | 6 | 42 | 66 | CP026001 |
| *Stenotrophomonas* genospecies 1<br>(Smc1; Sgn1) ZAC14D1 NAIMI4 6 | 4,700,343 | 4,128 | 6 | 45 | 63 | CP026000 |
| *Stenotrophomonas* genospecies 2<br>(Smc2; Sgn2) SAU14A NAIMI4 8 | 4,479,100 | 3,893 | 5 | 54 | 69 | CP025999 |
| *Stenotrophomonas* genospecies 2<br>(Smc2; Sgn2) YAU14A MKIMI4 1 | 4,487,007 | 3,918 | 7 | 43 | 67 | CP025998 |
| *Stenotrophomonas* genospecies 2<br>(Smc2; Sgn2) YAU14D1 LEIMI4 1 | 4,319,112 | 3,819 | 6 | 51 | 66 | CP025997 |
| *Stenotrophomonas* genospecies 2<br>(Smc2; Sgn2) ZAC14D2 NAIMI4 6 | 4,431,104 | 3,882 | 6 | 52 | 66 | CP025996 |
| *Stenotrophomonas* genospecies 2<br>(Smc2; Sgn2) ZAC14D2 NAIMI4 7 | 4,468,731 | 3,918 | 6 | 66 | 62 | CP025995 |

### Reference genomes

On August 1^st^, 2017, a total of 169 annotated *Stenotrophomonas* genome sequences were available in RefSeq, 134 of which were labeled as *S. maltophilia*. The corresponding GenBank files were retrieved, as well as the corresponding table with assembly metadata. Seven complete *Xanthomonas* spp. genomes were also downloaded to use them as outgroup sequences. In January 2018, the genome sequence of *S. bentonitica* strain VV6 was added to RefSeq and included in the revised version of this work to increase the taxon sampling.

### Computing consensus core- and pan-genomes with GET_HOMOLOGUES

We used GET_HOMOLOGUES (v05022018) (Contreras-Moreira and Vinuesa, 2013) to compute clusters of homologous gene families from the input genome sequences, as previously detailed (Vinuesa and Contreras-Moreira, 2015). Briefly, the source GenBank-formatted files were passed to get_homologues.pl and instructed to compute homologous gene clusters by running either our heuristic (fast) implementation of the bidirectional best-hit (BDBH) algorithm (’-b’) to explore the complete dataset, or the full BDBH, Clusters of Orthologous Groups - triangles (COGtriangles), and OrthoMCL (Markov Clustering of orthologues, OMCL) algorithms for the different sets of selected genomes, as detailed in the relevant sections and explained in the GET_HOMOLOGUES’s online manual (eead-csic-compbio.github.io/get_homologues/manual/manual.html). PFAM-domain scanning was enabled for the latter runs (-D flag). BLASTP hits were filtered by imposing a minimum of 90% alignment coverage (-C 90). The directories holding the results from the different runs were then passed to the auxiliary script compare_clusters.pl to compute either the consensus core genome (-t number_of_genomes) or pan-genome clusters (-t 0). The commands to achieve this can be found in the online tutorial https://vinuesa.github.io/get_phylomarkers/#get_homologues-get_phylomarkers-tutorials provided with the distribution.

### Overview of the computational steps performed by the GET_PHYLOMARKERS pipeline

Figure 1 presents a flow-chart that summarizes the computational steps performed by the pipeline, which are briefly described below. For an in-depth description of each step and associated parameters, as well as for a full version of the pipeline’s flow-chart, the reader is referred to the online manual (https://vinuesa.github.io/get_phylomarkers/). The pipeline is primarily intended to run DNA-based phylogenies (’-R 1 -t DNA’) on a collection of genomes from different species of the same genus or family. However, it can also select optimal markers for population genetics (’-R 2 -t DNA’), when the source genomes belong to the same species (not shown here). For more divergent genome sequences the pipeline should be run using protein sequences (’-R 1 -t PROT’). The analyses are started from the directory holding single-copy core-genome clusters generated either by ‘get_homologues.pl -e -t number_of_genomes’ or by ‘compare_clusters.pl -t number_of_genomes’. Note that both the protein (faa) and nucleotide (fna) FASTA files for the clusters are required, as detailed in the online tutorial (https://vinuesa.github.io/get_phylomarkers/#get_homologues-get_phylomarkers-tutorials). The former are first aligned with clustal-omega (Sievers et al., 2012) and then used by pal2nal (Suyama et al., 2006) to generate codon alignments. These are subsequently scanned with the Phi-test (Bruen et al., 2005) to identify and discard those with significant evidence for recombinant sequences. Maximum-likelihood phylogenies are inferred for each of the non-recombinant alignments using by default IQ-TREE v.1.6.2 (Nguyen et al., 2015), which will perform model selection with ModelFinder (Kalyaanamoorthy et al., 2017) using a subset of models and the ‘-fast’ flag enabled for rapid computation, as detailed in the online manual. Alternatively, FastTree v2.1.10 (Price et al., 2010) can be executed using the ‘-A F’ option, which will estimate phylogenies under the GTR+Gamma model. FastTree was compiled with double-precision enabled for maximum accuracy (see the manual for details). The resulting gene trees are screened to detect ‘outliers’ with help of the R package kdetrees (v.0.1.5) (Weyenberg et al., 2014, 2017). It implements a non-parametric test based on the distribution of tree topologies and branch lengths expected under the multispecies coalescent, identifying those phylogenies with unusual topologies or branch lengths. The stringency of the test can be controlled with the -*k* parameter (inter-quartile range multiplier for outlier detection, by default set to the standard 1.5). In a third step, the phylogenetic signal of each gene-tree is computed based on mean branch support values (Vinuesa et al., 2008), keeping only those above a user-defined mean Shimodaira-Hasegawa-like (SH-alrt) bipartition support (Anisimova and Gascuel, 2006) threshold (’-m 0.75’ by default). To make all the previous steps as fast as possible, they are run in parallel on multi-core machines using GNU parallel (Tange, 2011). The set of alignments passing all filters are concatenated and subjected to maximum-likelihood (ML) tree searching using by default IQ-TREE with model fitting to estimate the genomic species-tree.

**Figure 1.**
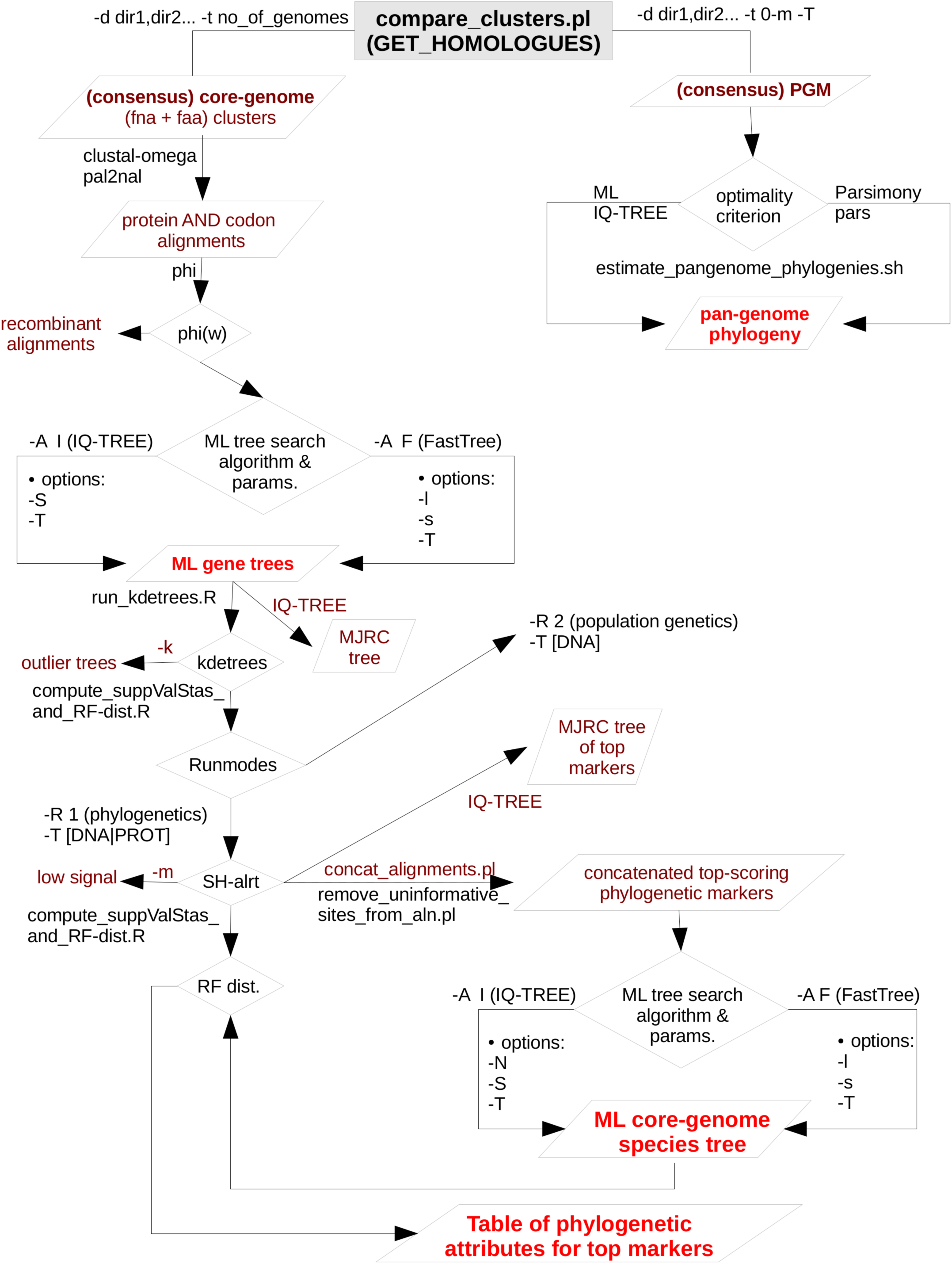
Simplified flow-chart of the GET_PHYLOMARKERS pipeline showing only those parts used and described in this work. The left branch, starting at the top of the diagram, is fully under control of the master script run_get_phylomarkes_pipeline.sh. The names of the worker scripts called by the master program are indicated on the relevant points along the flow. Steps involving repetitive computational processes, like generating multiple sequence alignments or inferring the corresponding gene trees, are run in parallel with the aid of GNU parallel, which is called from run_parallel_cmmds.pl. The right-hand branch at the top of the diagram summarizes the analyses that can be performed on the pan-genome matrix (PGM). In this work we only present the estimation of maximum-likelihood and parsimony pan-genome phylogenies. However, unsupervised learning approaches are provided by the hcluster_pangenome_matrix.sh script (not shown) for statistical analysis of the PGM. In addition, the plot_matrix_heatmap.sh script was used to analyze average nucleotide identity matrices generated by get_homologues.pl. It implements the unsupervised learning method described in this work to define the optimal number of clusters in such matrices. The plot_matrix_heatmap.sh script is distributed with the GET_HOMOLOGUES suite.

The complete GET_PHYLOMARKERS pipeline is launched with the master script run_get_phylomarkers_pipeline.sh, which calls a subset of auxiliary Bash, Perl and R programs to perform specific tasks. This architecture allows the user to run the individual steps separately, which adds convenient flexibility for advanced users (examples provided in the Supplementary Materials). The pipeline is highly customizable, and the reader is referred to the latest version of the online manual for the details of each option. However, the default values should produce satisfactory results for most purposes, as these were carefully selected based on the benchmark analysis presented in this work. All the source code is freely available under the GNU GENERAL PUBLIC LICENSE V3 from https://github.com/vinuesa/get_phylomarkers. Detailed installation instructions are provided (https://github.com/vinuesa/get_phylomarkers/blob/master/INSTALL.md), along with a hands-on tutorial (https://vinuesa.github.io/get_phylomarkers/). The software has been extensively tested on diverse Linux distributions (CentOS, Ubuntu and RedHat). In addition, a docker image bundling GET_HOMOLOGUES and GET_PHYLOMARKERS is available at https://hub.docker.com/r/csicunam/get_homologues/. We recommend running the docker image to avoid potential trouble with the installation and configuration of diverse dependencies (second party binaries, as well as Perl and R packages), making it easy to install on any architecture, including Windows, and to reproduce analyses with exactly the same software.

### Estimating maximum likelihood and parsimony pan-genome trees from the pan-genome matrix (PGM)

The GET_PHYLOMARKERS package contains auxiliary scripts to perform diverse clustering and phylogenetic analyses based on the pangenome_matrix_t0.* files returned by the compare_clusters.pl script (options ‘-t 0 -m’) from the GET_HOMOLOGUES suite. In this work, consensus PGMs (Vinuesa and Contreras-Moreira, 2015) were computed as explained in the online tutorial (https://vinuesa.github.io/get_phylomarkers/#get_homologues-get_phylomarkers-tutorials). These represent the intersection of the clusters generated by the COGtriangles and OMCL algorithms. Adding the -T flag to the previous command instructs compare_clusters.pl to compute a Wagner (multistate) parsimony tree from the pan-genome matrix, launching a tree search with 50 taxon jumbles with pars from the PHYLIP (Felsenstein, 2004b) package (v.3.69). A more thorough and customized ML or parsimony analysis of the PGM can be performed with the aid of the auxiliary script estimate_pangenome_phylogenies.sh, bundled with GET_PHYLOMARKERS. By default this script performs a ML tree-search using IQ-TREE v1.6.2 (Nguyen et al., 2015). It will first call ModelFinder (Kalyaanamoorthy et al., 2017) using the JC2 and GTR2 base models for binary data, the latter accounting for unequal state frequencies. The best fitting base model + ascertain bias correction + among-site rate variation parameters are selected using the Akaike Information Criterion (AIC). IQ-TREE (Nguyen et al., 2015) is then called to perform a ML tree search under the selected model with branch support estimation. These are estimated using approximate Bayesian posterior probabilities (aBypp), a popular single branch test (Guindon et al., 2010), as well as the recently developed ultrafast-bootstrap2 (UFBoot2) test (Hoang et al., 2017). In addition, the user may choose to run a parsimony analysis with bootstrapping on the PGM, as detailed in the online manual and illustrated in the tutorial. Note however, that the parsimony search with bootstrapping is much slower than the default ML search.

### Unsupervised learning methods for the analysis of pairwise average nucleotide (ANI) and aminoacid (AAI) identity matrices

The GET_HOMOLOGUES distribution contains the plot_matrix_heatmap.sh script which generates ordered heatmaps with attached row and column dendrograms from squared tab-separated numeric matrices. These can be presence/absence PGM matrices or similarity / identity matrices, as those produced with the get_homologues -A option. Optionally, the input cgANIb matrix can be converted to a distance matrix to compute a neighbor joining tree, which makes the visualization of relationships in large ANI matrices easier. Recently added functionality includes reducing excessive redundancy in the tab-delimited ANI matrix file (-c max_identity_cut-off_value) and sub-setting the matrix with regular expressions, to focus the analysis on particular genomes extracted from the full cgANIb matrix. From version 1.0 onwards, the mean silhouette-width (Rousseeuw, 1987) goodness of clustering statistics to determine the optimal number of clusters automatically. The script currently depends on the R packages ape (Popescu et al., 2012), dendextend (https://cran.r-project.org/package=dendextend), factoextra (https://cran.r-project.org/package=factoextra) and gplots (https://CRAN.R-project.org/package=gplots).

## RESULTS

### Ten new complete genome assemblies for the Mexican environmental *Stenotrophomonas maltophilia* complex isolates previously classified as genospecies 1 (Smc1) and 2 (Smc2)

In this study we report the sequencing and assembly of five isolates each from the genospecies 1 (Smc1) and 2 (Smc2) recovered from rivers in Central Mexico, previously reported in our extensive MLSA study of the genus *Stenotrophomonas* (Ochoa-Sánchez and Vinuesa, 2017). All assemblies resulted in a single chromosome with gaps. No plasmids were detected. A summary of the annotated features for each genome are presented in Table 1. Assembly details for each genome are provided in supplementary Table S1.

### Rapid phylogenetic exploration of *Stenotrophomonas* genome sequences available at NCBI’s RefSeq repository running GET_PHYLOMARKERS in fast runmode

A total of 170 *Stenotrophomonas* and 7 *Xanthomona*s reference genomes were retrieved from RefSeq (see methods). Figure 2A depicts parallel density plots showing the distribution of the number of fragments for the *Stenotrophomonas* assemblies at the Complete (*n* = 16), Chromosome (*n* = 3), Scaffold (*n* = 63) and Contig (*n* = 88) finishing levels. The distributions have conspicuous long tails, with an overall mean and median number of fragments of ~238 and ~163, respectively. The table insets in Fig. 2A provide additional descriptive statistics of the distributions. A first GET_HOMOLOGUES run was launched using this dataset (*n* = 177) with two objectives: *i*) to test its performance with a relatively large set of genomes and *ii*) to get an overview of their evolutionary relationships to select a non-redundant set of those with the best assemblies. For this analysis, GET_HOMOLOGUES was run in its “fast-BDBH” mode (-b), on 60 cores (-n 60; AMD Opteron^TM^ Processor 6380, 2500.155 MHz), and imposing a stringent 90% coverage cut-off for BLASTP alignments (-C 90), excluding inparalogues (-e). This analysis took 1h:32m:13s to complete and identified 132 core genes. These were fed into the GET_PHYLOMARKERS pipeline, which was executed using a default FastTree search with the following command line: run_get_phylomarkers_pipeline.sh -R 1 -t DNA -A F, which took 8m:1s to complete on the same number of cores. Only 79 alignments passed the Phi recombination test. Thirteen of them failed to pass the downstream kdetree test. The phylogenetic signal test excluded nine additional loci with average SH-alrt values < 0.70. Only 57 alignments passed all filters and were concatenated into a supermatrix of 38,415 aligned residues, which were collapsed to 19,129 non-gapped and variable sites. A standard FastTree maximum-likelihood tree-search was launched with the command: ‘run_get_phylomarkers_pipeline.sh -R 1 -t DNA -A F’. The resulting phylogeny (ln*L* = −475237.540) is shown in supplementary Figure S1. Based on this tree and the level of assembly completeness for each genome (Fig. 2A), we decided to discard those with > 300 contigs (Fig. 2B). This resulted in the loss of 19 genomes labeled as *S. maltophilia.* However, we retained *S. pictorum* JCM 9942, a highly fragmented genome with 829 contigs (Patil et al., 2016) to maximize taxon sampling. Several *S. maltophilia* subclades contained identical sequences (Fig. S1) and were trimmed, retaining only the assembly with the lowest numbers of scaffolds or contigs.

**Figure 2.**
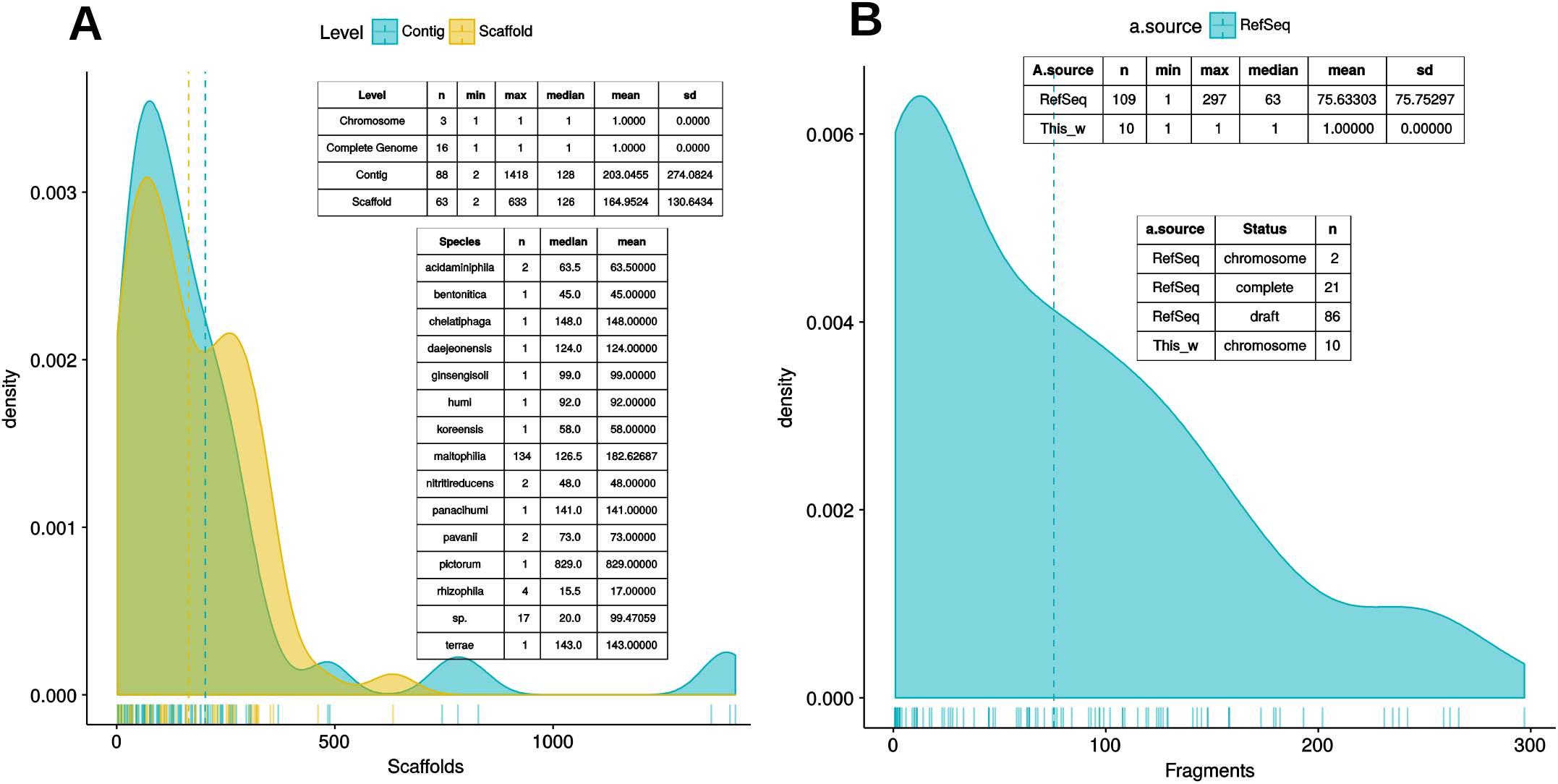
Density plots showing the distribution of the number of fragments of the *Stenotrophomonas* genomes available in RefSeq as of August 2017, plus the genome of *S. bentonitica* VV6, released in January 2018. **A**) Distribution of the number of fragments in the assemblies of 170 annotated *Stenotrophomonas* genomes as a function of assembly status (contigs vs. scaffolds) plus 7 *Xanthomonas* genomes used as outgroup to root the tree. Inset tables provide additional summary statistics of the RefSeq assemblies. **B**) Distribution of the number of fragments in the assemblies of the 119 genomes selected for the analyses presented in this study, which include 102 reference *Stenotrophomonas* genomes, 10 new genomes generated for this study, and 7 complete *Xanthomonas* spp. genomes.

### Selection of a stringently defined set of orthologous genes using GET_HOMOLOGUES

After the quality and redundancy filtering described in the previous section, 109 reference genomes (102 *Stenotrophomonas* + 7 *Xanthomonas*) were retained for more detailed investigation. Table S2 provides an overview of them. To this set we added the 10 new genomes reported in this study (Table 1). Figure 2B depicts parallel density plots summarizing the distribution of number of contigs/scaffolds in the selected reference genomes and the new genomes for the Mexican environmental Smc isolates previously classified as genospecies 1 (Smc1) and 2 (Smc2) (Ochoa-Sánchez and Vinuesa, 2017). A high stringency consensus core-genome containing 239 gene families was computed as the intersection of the clusters generated by the BDBH, COG-triangles and OMCL algorithms (Fig 3A).

**Figure 3.**
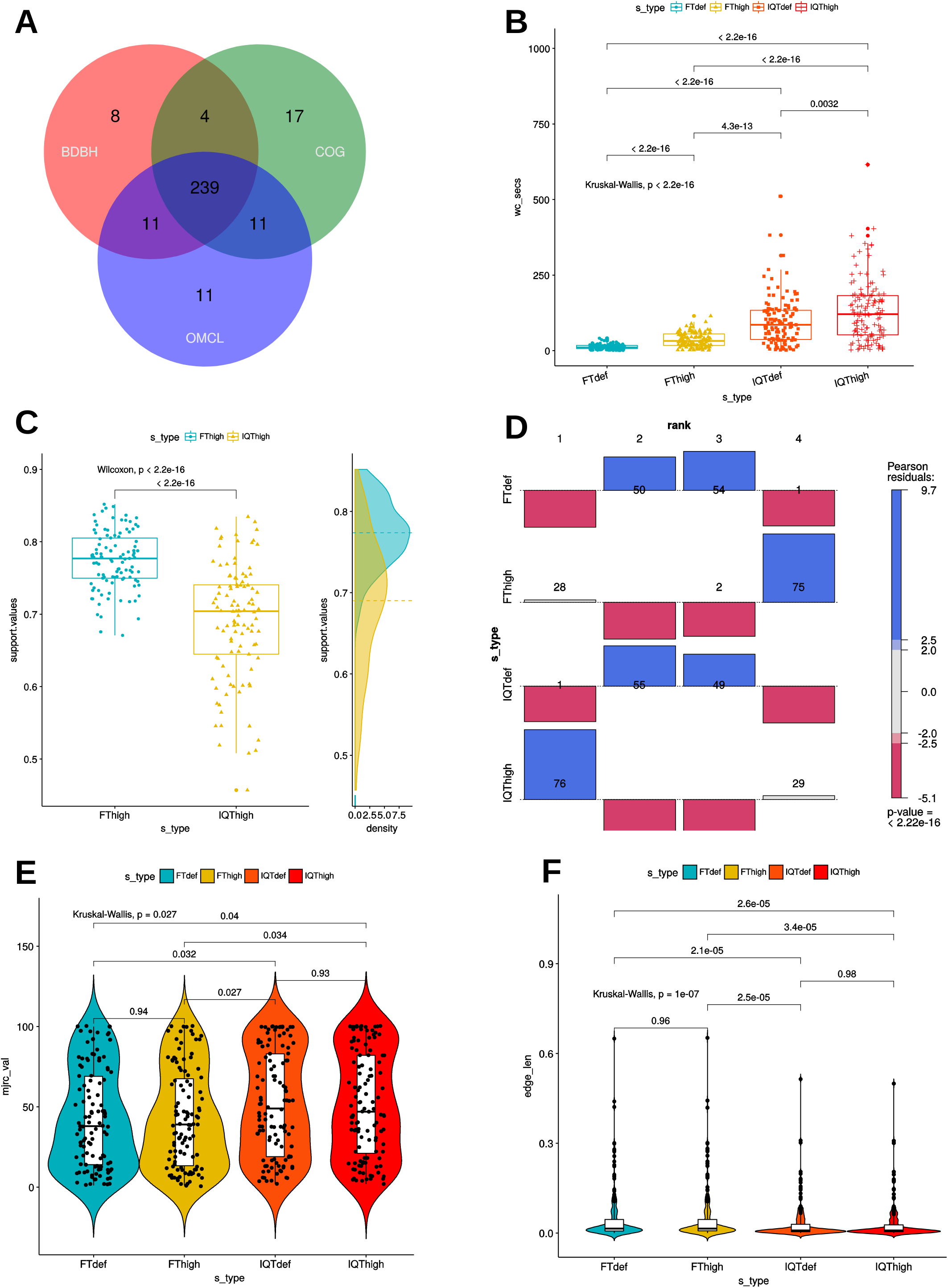
Combined filtering actions performed by GET_HOMOLOGUES and GET_PHYLOMARKERS to select top-ranking phylogenetic markers to be concatenated for phylogenomic analyses, and benchmark results of the performance of the FastTree (FT) and IQ-TREE (IQT) maximum-likelihood (ML) phylogeny inference programs. **A**) Venn-diagram indicating the number consensus and algorithm-specific core-genome orthologous clusters. **B**) Parallel box-plots summarizing the computation time required by FT and IQT when run under “default” (FTdef, IQTdef) and thorough (FThigh, IQThigh) search modes (s_type) on the 239 consensus clusters, as detailed in the main text. Statistical significance of differences between treatments were computed with the Kruskal-Wallis (robust, non-parametric, ANOVA-like) test. **C**) Distribution of SH-alrt branch support values of gene-trees found by the FThigh and IQThigh searches. Statistical significance of differences between the paired samples was computed with the Wilcoxon signed-rank test. This is a non-parametric alternative to paired t-test used to compare paired data when they are not normally distributed. **D**) Association plot (computed with the vcd package) summarizing the results of multi-way Chi-Square analyses of the ln*L* score ranks (1 to 4, meaning best to worst) of the ML gene-trees computed from the set of 105 codon alignments passing the kdetrees filter in the IQThigh run (**Table 2**) for each search-type. The height and color-shading of the bars indicate the magnitude and significance level of the Pearson residuals. **E**) Statistical analysis (Kruskal-Wallis test) of the distribution of consensus values from majority-rule consensus trees computed from the gene trees passing all the filters, as a function of search-type. **D**) Statistical analysis (Kruskal-Wallis test) of the distribution of the edge-lengths of species-trees computed from the concatenated top-scoring markers, as a function of search-type.

### GET_PHYLOMARKERS in action: benchmarking the performance of FastTree and IQ-TREE to select top-scoring markers for phylogenomics

The set of 239 consensus core-genome clusters (Fig. 3A) was used to launch multiple instances of the GET_PHYLOMARKERS pipeline to evaluate the phylogenetic performance of FastTree (FT; v2.1.10) and IQ-TREE (IQT; v1.6.2), two popular fast maximum-likelihood (ML) tree searching algorithms. Our benchmark was designed to compare: *i*) the execution times of the FT vs. IQT runs under default (FTdef, IQTdef) and thorough (FThigh, IQThigh) search modes (see methods and online manual for their parameterization details); *ii*) the phylogenetic resolution (average support values) of gene trees estimated by FT and IQT under both search modes; *iii*) the rank of ln*L* scores of the gene trees found in those searches for each locus; *iv*) the distribution of consensus values of each node in majority rule consensus trees computed from the gene trees found by each search type; *v*) the distribution of edge-lengths in the species-trees computed by each search type. The results of these analyses are summarized in Table 2 and in Figure 3. The first steps of the pipeline (Fig. 1) comprise the generation of codon alignments and their analysis to identify potential recombination events. Only 127 alignments (53.14 %) passed the Phi-test (Table 2). Phylogenetic analyses start downstream of the recombination test (Fig. 1). The computation times required by the two algorithms and search intensity levels were significantly different (Kruskal-Wallis, *p* < 2.2e-16), FastTree being always the fastest, and displaying the lowest dispersion of compute times across trees (Fig. 3B). This is not surprising, as IQT searches involved selecting the best substitution model among a range of base models (see methods and online manual) and fitting additional parameters (+G+ASC+I+F+R) to account for heterogeneous base frequencies and rate-variation across sites. In contrast, FT searches just estimated the parameter values for the general time-reversible (GTR) model, and among-site rate variation was modeled fitting a gamma distribution with 20 rate categories (+G), as summarized in Table 2. Similar numbers of “outlier” trees (range 18:22) were detected by the kdetrees-test in the four search types (Table 2). However, the distributions of SH-alrt support values are strikingly different for both search algorithms (Wilcoxon, *p* < 2.2e-16), revealing that gene-trees found by IQT have a much lower average support than those found by FT (Fig 3C). Consequently, the former searches were significantly more efficient to identify gene trees with low average branch support values (Table 2 and Fig. 3C). This result is in line with the well-established fact that poorly fitting and under-parameterized models produce less reliable tree branch lengths and overestimate branch support (Posada and Buckley, 2004), implying that the FT phylogenies may suffer from clade over-credibility. These results demonstrate that: *i*) FT-based searches are significantly faster than those performed with IQT, and *ii*) that IQT has a significantly higher discrimination power for phylogenetic signal than FT. Due to the fact that the number of top-scoring alignments selected by the two algorithms for concatenation is notably different (Table 2), the ln*L* scores of the resulting species-trees are not comparable (Table 2). Therefore, in order to further evaluate the quality of the gene-trees found by the four search strategies, we performed an additional benchmark under highly standardized conditions, based on the 105 optimal alignments that passed the kdetrees-test in the IQThigh search (Table 2). Gene trees were estimated for each of these alignments using the four search strategies (FTdef, IQTdef, FThigh and IQThigh) and their ln*L* scores ranked for each gene tree. An association analysis (deviation from independence in a multi-way chi-squared test) was performed on the ln*L* ranks (1 to 4, coding for highest to lowest ln*L* scores, respectively) attained by each search type for each gene tree. As shown in Fig. 3D, the IQThigh search was the winner, attaining the first rank (highest ln*L* score) in 76/105 of the searches (72.38%), way ahead of the number of FThigh (26%), and IQTdef (0.009%) searches that ranked in the first position (highest ln*L* score for a particular alignment). A similar analysis performed on the full set of input alignments (*n* = 239) indicated that when operating on an unfiltered set, the difference in performance was even more striking, with IQT-based searches occupying > 97 % of the first rank positions (data not shown). These results highlight two points: *i*) the importance of proper model selection and thorough tree searching in phylogenetic inference and *ii*) that IQT generally finds better trees than FT. Finally, we evaluated additional phylogenetic attributes of the species-trees computed by each search type, either as the majority rule consensus (mjrc) tree of top-scoring gene-trees, or as the tree estimated from the supermatrices of concatenated alignments. Figure 3E shows the distribution of mjrc values of the mjrc trees computed by each search type, which can be interpreted as a proxy for the level phylogenetic congruence among the source trees. These values were significantly higher for the IQT than in the FT searches (Kruskal-Wallis, *p* = 0.027), with a higher number of 100% mjrc clusters found in the former than in the latter type of trees (Fig. 3E). An analysis of the distribution of edge-lengths of the species-trees inferred from the concatenated alignments revealed that those found in IQT searches had significantly (Kruskal-Wallis, *p* = 1e-07) shorter edges (branches) than those estimated by FT (Fig. 3F). This highlights again the importance of adequate substitution models for proper edge-length estimation. Tree-lengths (sum of edge lengths) of the species-trees found in IQT-based searches are about 0.63 times shorter than those found by FT (Fig. S2). As a final exercise, we computed the Robinson-Foulds (RF) distances of each gene tree found in a given search type to the species tree inferred from the corresponding supermatrix. The most striking result of this analysis was that no single gene-tree had the same topology as the species tree inferred from the concatenated top-scoring alignments (Fig. S3).

**Table 2.**
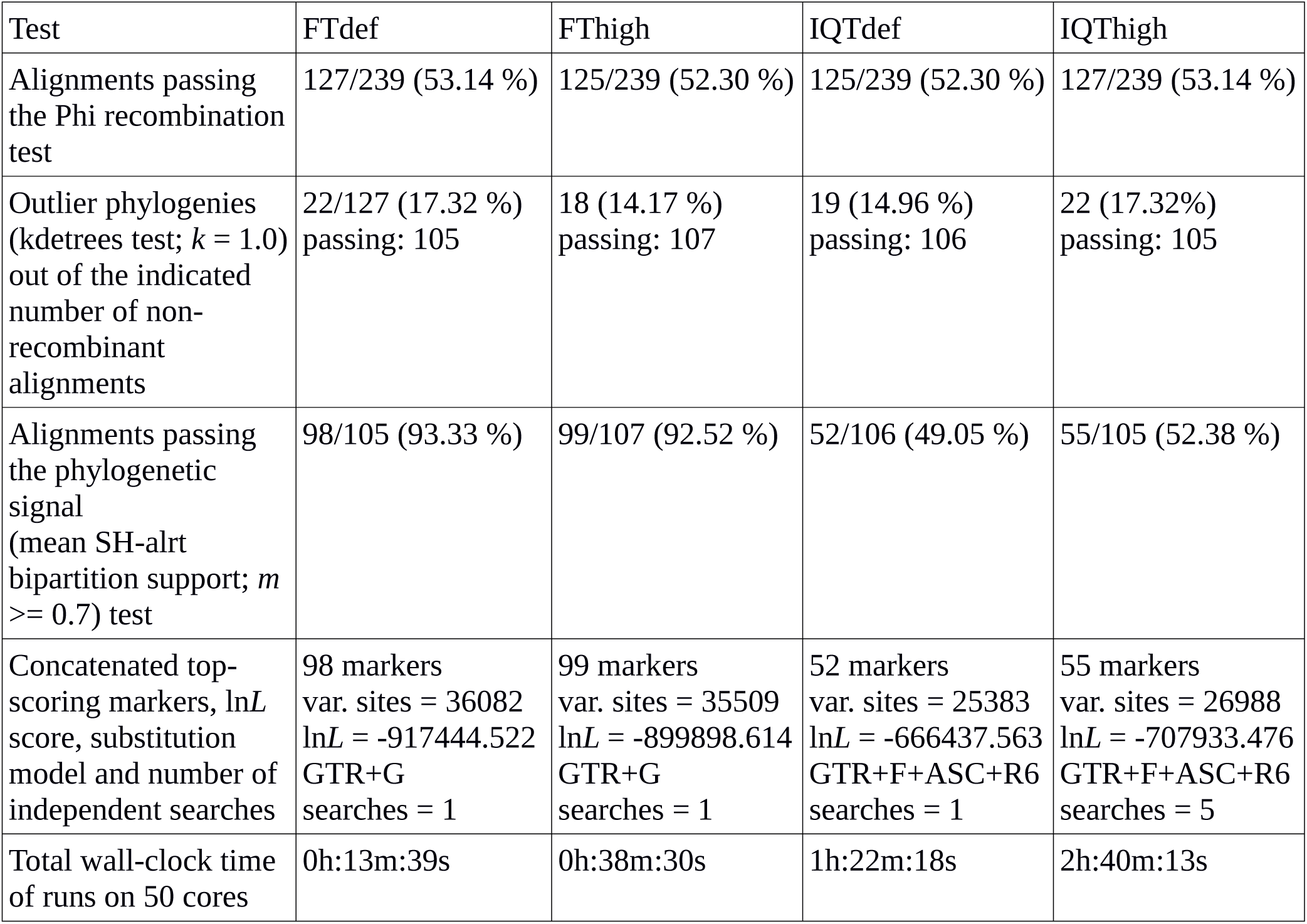
Comparative benchmark analysis of the filtering performance of the GET_PHYLOMARKERS pipeline when run using the FastTree (FT) and IQ-TREE (IQT) maximum-likelihood algorithms under default and high search-intensity levels. The analyses were started with the stringently defined set of 239 consensus core-genome clusters computed by GET_HOMOLOGUES for a dataset of 119 genomes (112 *Stenotrophomonas* spp. and 7 *Xanthomonas* spp.).

### Effect of tree-search intensity on the quality of the species trees found by IQT-REE and FastTree

Given the astronomical number of different topologies that exist for 119 terminals, we decided to evaluate the effect of tree-search thoroughness on the quality of the trees found by FT and IQT, measured as their log-likelihood (ln*L*) score. To make the results comparable across search algorithms, we used the supermatrix of 55 top-scoring markers (25,896 variable, non-gapped sites) selected by the IQThigh run (Table 2). One thousand FT searches were launched from the same number of random topologies computed with the aid of a custom Perl script. In addition, a standard FT search was started from the default BioNJ tree. All these searches were run in “thorough” mode (-quiet -nt -gtr -bionj -slow -slownni -gamma -mlacc 3 -spr 16 -sprlength 10) on 50 cores. The resulting ln*L* profile for this search is presented in Figure 4A, which reached a maximal score of −717195.373. This is 121.281 ln*L* units better than the score of the best tree found in the search started from the BioNJ seed tree (ln*L* −717316.654, lower discontinuous blue line). In addition, 50 independent tree searches were run with IQ-TREE under the best fitting model previously found (Table 2), using the shell loop command (# 5) provided in the Supplementary Material. The corresponding ln*L* profile of this search is shown in Fig. 4B, which found a maximum-scoring tree with a score of −707932.468. This is only 8.105 ln*L* units better than the worst tree found in that same search (Fig. 4B). Importantly, the best tree found in the IQT-search is 9262.905 ln*L* units better that of the best tree found in the FT search, despite the much higher number of seed trees used for the latter. This result clearly demonstrates the superiority of the IQ-TREE algorithm for ML tree searching. Based on this evidence, and that presented in the previous section (Table 2; Fig. 3), IQ-TREE was chosen as the default tree-search algorithm used by GET_PHYLOMARKERS. The Robinson-Foulds distance between both trees was 46.

### A robust genomic species phylogeny for the genus *Stenotrophomonas*: taxonomic implications and identification of multiple misclassified genomes

Figure 5 displays the best ML phylogeny found in the IQ-TREE search (Fig. 4B) described in the previous section. This is a highly resolved phylogeny. All bipartitions have an approximate Bayesian posterior probability (aBypp) *p* >= 0.95. It was rooted at the branch subtending the *Xanthomonas* spp. clade, used as an outgroup. A first taxonomic inconsistency revealed by this phylogeny is the placement of *S. panacihumi* within the latter clade, making the genus *Stenotrophomonas* paraphyletic. It is worth noting that *S. panacihumi* is a non-validly described, and poorly characterized species (Yi et al., 2010). The genus *Stenotrophomonas*, as currently defined, and excluding *S. panacihumi*, consists of two major clades, labeled as I and II in Fig. 5, as previously defined (Ochoa-Sánchez and Vinuesa, 2017). Clade I groups environmental isolates, recovered from different ecosystems, mostly soils and plant surfaces, classified as *S. ginsengisoli* (Kim et al., 2010), *S. koreensis* (Yang et al., 2006), *S. daejeonensis* (Lee et al., 2011), *S. nitritireducens* (Finkmann et al., 2000), *S. acidaminiphila* (Labat et al., 2002), *S. humi* and *S. terrae* (Heylen et al., 2007). The recently described *S. pictorum* (Ouattara et al., 2017) is also included in clade I. These are all rather poorly studied species, for which only one or a few strains have been considered in the corresponding species description or to study particular aspects of their biology. None of these species have been reported as opportunistic pathogens, but some contain promising strains for plant growth-promotion and bio-remediation. Particularly notorious are the disproportionally long terminal branches (heterotachy) of *S. ginsengisoli* and *S. koreensis* (Fig. 5). The potential distortion of these long branches on the estimated phylogeny needs to be evaluated in future work.

**Figure 4.**
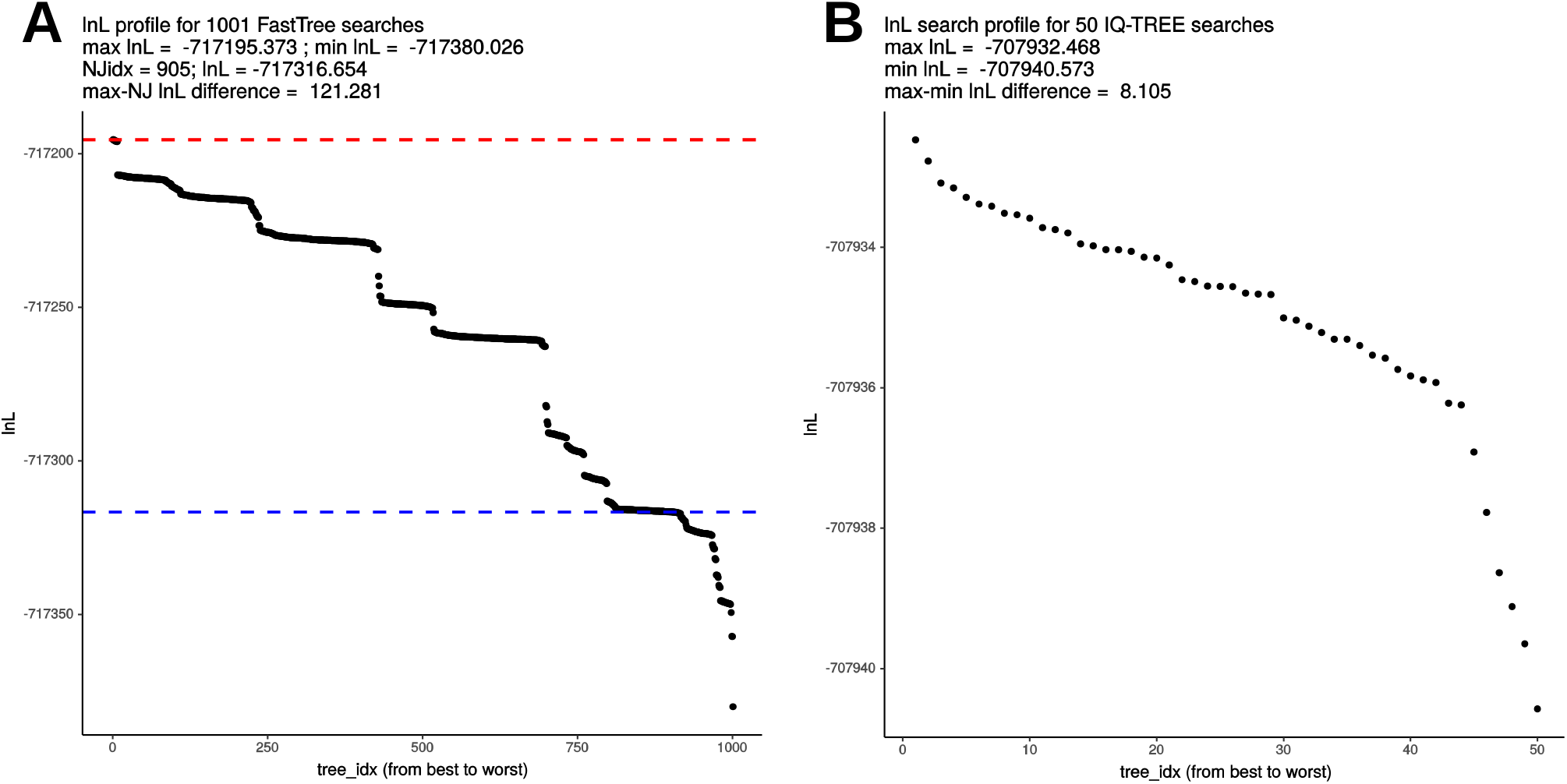
Comparative analysis of log-likelihood tree search profiles. **A**) Sorted ln*L* profile of FastTree (FT) tree searches launched from 1000 random trees + 1 BioNJ phylogeny, using the “thorough” tree-search settings described in the main text and the 55 top-ranking markers (26,988 non-gapped, variable sites) selected by the IQThigh run for 119 genomes (**Table 2**). The dashed blue line indicates the score of the search initiated from the BioNJ tree. **B**) Sorted ln*L* profile of 50 independently launched IQ-TREE (IQT) searches under the best-fitting model using the same matrix as for the FT search.

**Figure 5.**
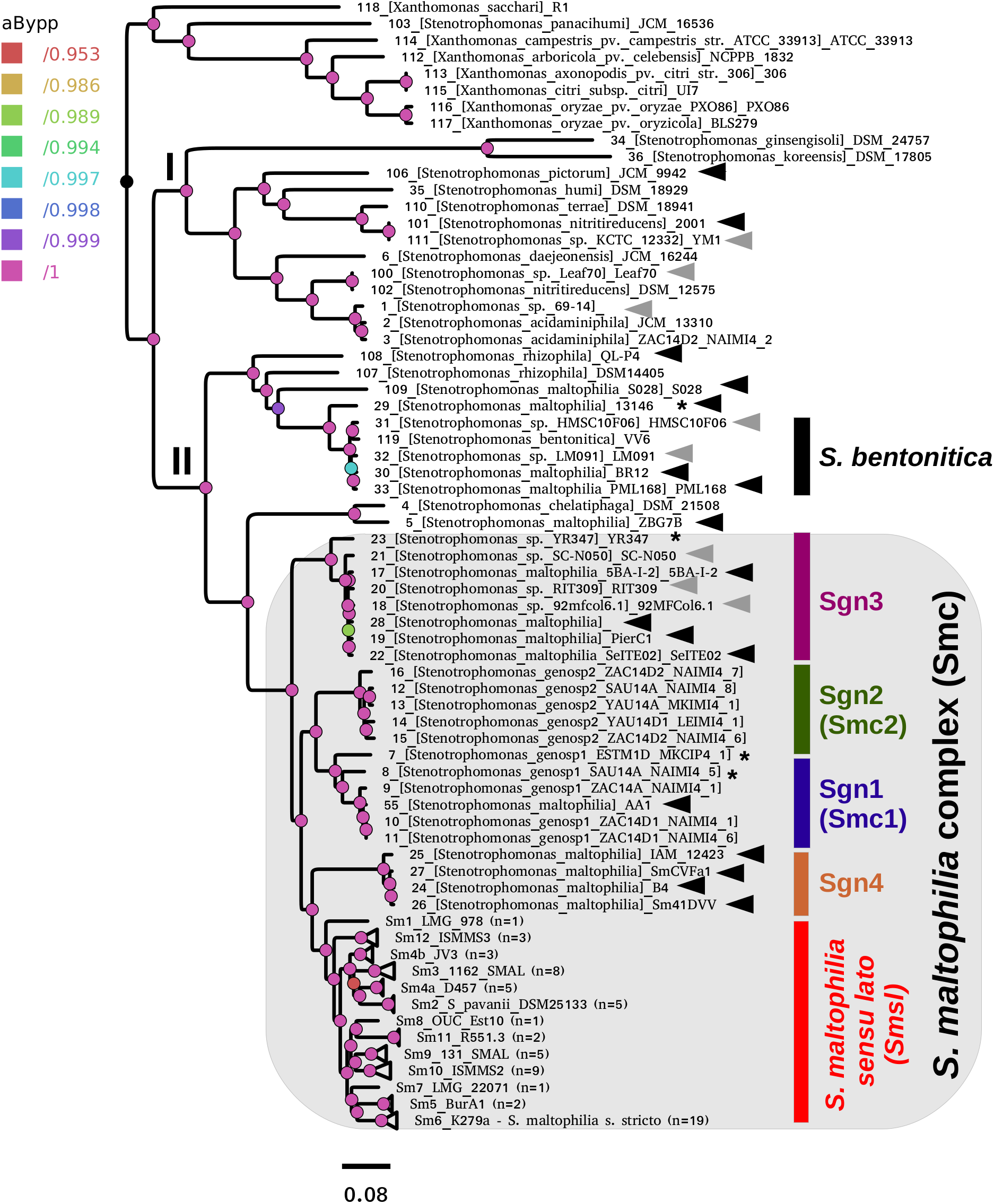
Best maximum-likelihood core-genome phylogeny for the genus *Stenotrophomonas* found in the IQ-TREE search described in **Fig. 4B**, based on the supermatrix obtained by concatenation of 55 top-ranking alignments (**Table 2**). The tree was rooted using the *Xanthomonas* spp. sequences as the outgroup. Arrows highlight genomes not grouping in the *S. maltophilia sensu lato* clade (Smsl), for which we suggest a reclassification, as summarized in **Table 3**. Black arrows indicate misclassified strains, while gray ones mark unclassified genomes. The shaded area highlights the strains considered as members of the *S. maltophilia* complex (Smc). The genospecies 1 and 2 (Sgn1 = Smc1; Sgn2 = Smc2) were previously recognized as separate species-like lineages by Ochoa-Sánchez and Vinuesa (2017). Strains grouped in the Smsl clade are collapsed into sub-clades that are perfectly consistent with the cluster analysis of core-genome average nucleotide identity (cgANIb) values presented in **Fig. 7** at a cutoff-value of 95.9%. Integers in parentheses correspond to the number of genomes in each collapsed clade. **Supplementary Figure S4** displays the same tree in non-collapsed form. Strains from genospecies 1, 3 and 5 (Sgn1, Sgn3, Sgn5) marked with an asterisk may represent additional species, according to the cgANIb values. Nodes are colored according to the lateral scale, which indicates the approximate Bayesian posterior probability values. The scale bar represents the number of expected substitutions per site under the best-fitting GTR+ASC+F+R6 model.

Clade II contains the species *S. rhizophila* (Berg et al., 2002), *S. chelatiphaga* (Kaparullina et al., 2009), the recently described *S. bentonitica* (Sánchez-Castro et al., 2017), along with multiple species and genospecies lumped in the *S. maltophilia* complex (Smc; shaded area in Fig. 5) (Berg and Martinez, 2015; Svensson-Stadler et al., 2012). The Smc includes the validly described *S. maltophilia* (Palleroni and Bradbury, 1993) and *S. pavanii* (Ramos et al., 2011) (collapsed subclades Sm6 and Sm2, respectively, located within the clade labeled as *S. maltophilia sensu lato* in Figure 5), along with at least four undescribed genospecies (Sgn1-Sgn4) recently identified in our MLSA study of the genus (Ochoa-Sánchez and Vinuesa, 2017). In light of this phylogeny, we discovered 14 misclassified RefSeq genome sequences (out of 119; ~11.76 %), 12 of them labeled as *S. maltophilia*. These genomes are highlighted with black arrows in Figure 5. The phylogeny also supports the classification, either as a validly published species, or as new genospecies, of 8 (~ 6.72 %) additional RefSeq genomes (gray arrows) lacking a species assignation in the RefSeq record, as summarized in Table 3. In addition, the phylogeny resolved 13 highly supported lineages (aBypp > 0.95) within the *S. maltophilia sensu lato* (Smsl) cluster, shown as collapsed clades. They have a core-genome average nucleotide identity > 96 % (Fig. 5). These lineages may represent 11 additional species in the Smsl clade, as detailed in following sections. Supplementary Figure S4 shows the non-collapsed version of the species-tree displayed in Figure 5.

**Table 3.** RefSeq genome sequences reclassified in this study based on the diverse genomic evidence presented herein (see Figures 5, 6 and 7).

| DEFINITION (RefSeq classification) | Species/reclassification* | Status | Fragments | BioProject | BioSample | PMID |
| --- | --- | --- | --- | --- | --- | --- |
| *Stenotrophomonas* sp. 69-14 | *S. acidaminiphila* | draft | 27 | PRJNA279279 | SAMN05660631 | NA |
| *Stenotrophomonas maltophilia* ZBG7B | *S. chelatiphaga* | draft | 145 | PRJNA272355 | SAMN03280975 | 26659682 |
| *Stenotrophomonas maltophilia* AA1 | genospecies 2 (Sgn2/Smc2) | complete | 1 | PRJNA224116 | SAMN06130959 | 28275097 |
| *Stenotrophomonas maltophilia* 5BA-I-2 | genospecies 3 | draft | 4 | PRJNA224116 | SAMN02641498 | 24604648 |
| *Stenotrophomonas* sp. 92mfcol6.1 | genospecies 3 | draft | 11 | PRJNA224116 | SAMN04488690 | NA |
| *Stenotrophomonas maltophilia* PierC1 | genospecies 3 | draft | 59 | PRJEB8824 | SAMEA3309462 | 26276674 |
| *Stenotrophomonas* sp. RIT309 | genospecies 3 | draft | 45 | PRJNA224116 | SAMN02676627 | 24812212 |
| *Stenotrophomonas* sp. SC-N050 | genospecies 3 | draft | 24 | PRJNA224116 | SAMN05720615 | NA |
| *Stenotrophomonas maltophilia* SeITE02 | genospecies 3 | draft | 63 | PRJNA224116 | SAMEA3138997 | 24812214 |
| *Stenotrophomonas* sp. YR347 | genospecies 3 | draft | 11 | PRJNA224116 | SAMN05518671 | NA |
| *Stenotrophomonas maltophilia* B4 | genospecies 4 | draft | 180 | PRJNA224116 | SAMN03753636 | NA |
| *Stenotrophomonas maltophilia* Sm41DVV | genospecies 4 | draft | 26 | PRJNA323790 | SAMN05188789 | 27540065 |
| *Stenotrophomonas maltophilia* SmCVFa1 | genospecies 4 | draft | 30 | PRJNA323845 | SAMN05190067 | 27540065 |
| *Stenotrophomonas maltophilia* 13146 | *S. bentonitica* complex | draft | 60 | PRJNA224116 | SAMN07237143 | NA |
| *Stenotrophomonas maltophilia* BR12S | *S. bentonitica* | draft | 80 | PRJNA224116 | SAMN03456145 | 26472823 |
| *Stenotrophomonas* sp. HMSC10F07 | *S. bentonitica* | draft | 63 | PRJNA269850 | SAMN03287020 | NA |
| *Stenotrophomonas* sp. LM091 | *S. bentonitica* | complete | 1 | PRJNA344031 | SAMN05818440 | 27979933 |
| *Stenotrophomonas maltophilia* PML168 | *S. bentonitica* | draft | 97 | PRJNA224116 | SAMEA2272452 | 22887661 |
| *Stenotrophomonas* sp. Leaf70 | *S. nitritireducens* | draft | 11 | PRJNA224116 | SAMN04151613 | 26633631 |
| *Stenotrophomonas* sp. KCTC 12332 | *S. terrae* complex | complete | 1 | PRJNA310387 | SAMN04451766 | 28689013 |
| *Stenotrophomonas nitritireducens* 2001 | *S. terrae* complex | complete | 1 | PRJNA224116 | SAMN05428703 | NA |
| *Stenotrophomonas maltophilia* S028 | *Stenotrophomonas* sp. |  |  |  |  |  |
| *Stenotrophomonas rhizophila* QL-P4 | *Stenotrophomonas* sp. | complete | 1 | PRJNA326321 | SAMN05276013 | NA |

\*The numbered genospecies correspond to novel unnamed species identified by Ochoa-Sánchez and Vinuesa (2017) and in this study. Unnamed species classified as members of the *S. maltophilia sensu* *lato* clade (Fig. 5) are labeled as *S. maltophilia s. l.* Strains assigned to the *S. terrae* complex most likely represent novel species related to *S. terrae*.

No genome sequences, nor MLSA data are available for the recently described *S. tumulicola* (Handa et al., 2016).

### Pan-genome phylogenies for the genus *Stenotrophomonas* recover the same species clades as the core-genome phylogeny

A limitation of core-genome phylogenies is that they are estimated from the small fraction of single-copy genes shared by all organisms under study. Genes encoding adaptive traits relevant for niche-differentiation and subsequent speciation events typically display a lineage-specific distribution. Hence, phylogenetic analysis of pan-genomes, based on their differential gene-composition profiles, provide a complementary, more resolved and often illuminating perspective on the evolutionary relationships between species.

A consensus pan-genome matrix (PGM) containing 29,623 clusters was computed from the intersection of the clusters generated by the COG-triangles and OMCL algorithms (Figure 6). This PGM was subjected to ML tree searching using the binary and morphological models implemented in IQ-TREE for phylogenetic analysis of discrete characters with the aid of the estimate_pangenome_phylogenies.sh script bundled with GET_PHYLOMARKERS (Fig. 1). As shown in the tabular inset of Figure 6, the binary GTR2+FO+R4 model was by large the best-fitting one (with the smallest AIC and BIC values). Twenty five independent IQ-TREE searches were performed on the consensus PGM with the best-fitting model. The best tree found is presented in Figure 6, rooted with the *Xanthomonas* spp. outgroup sequences. It depicts the evolutionary relationships of the 119 genomes based on their gene content (presence-absence) profiles. The numbers on the nodes indicate the approximate Bayesian posterior probabilities (aBypp) / UFBoot2 support values (see methods). The same tree, but without collapsing clades, is presented in the supplementary figure S5. This phylogeny resolves exactly the same species-like clades highlighted on the core-genome phylogeny presented in Figure 5, which are also grouped in the two major clades I and II. These are labeled with the same names and color-codes, for easy cross-comparison. However, there are some notorious differences in the phylogenetic relationships between species on both trees, like the placement of *S. panacihumi* outside of the *Xanthomonas* clade, and the sister relation of genospecies 3 (Sgnp3) to the *S. maltophilia sensu lato* clade. These same relationships were found in a multi-state (Wagner) parsimony phylogeny of the PGM shown in Supplementary Figure S6. In summary, all core-genome and pan-genome analyses presented consistently support our previous claim that the five genospecies defined in our MLSA study represent distinct species and support the existence of multiple cryptic species within the Smsl clade, as defined in Figure 5.

**Figure 6.**
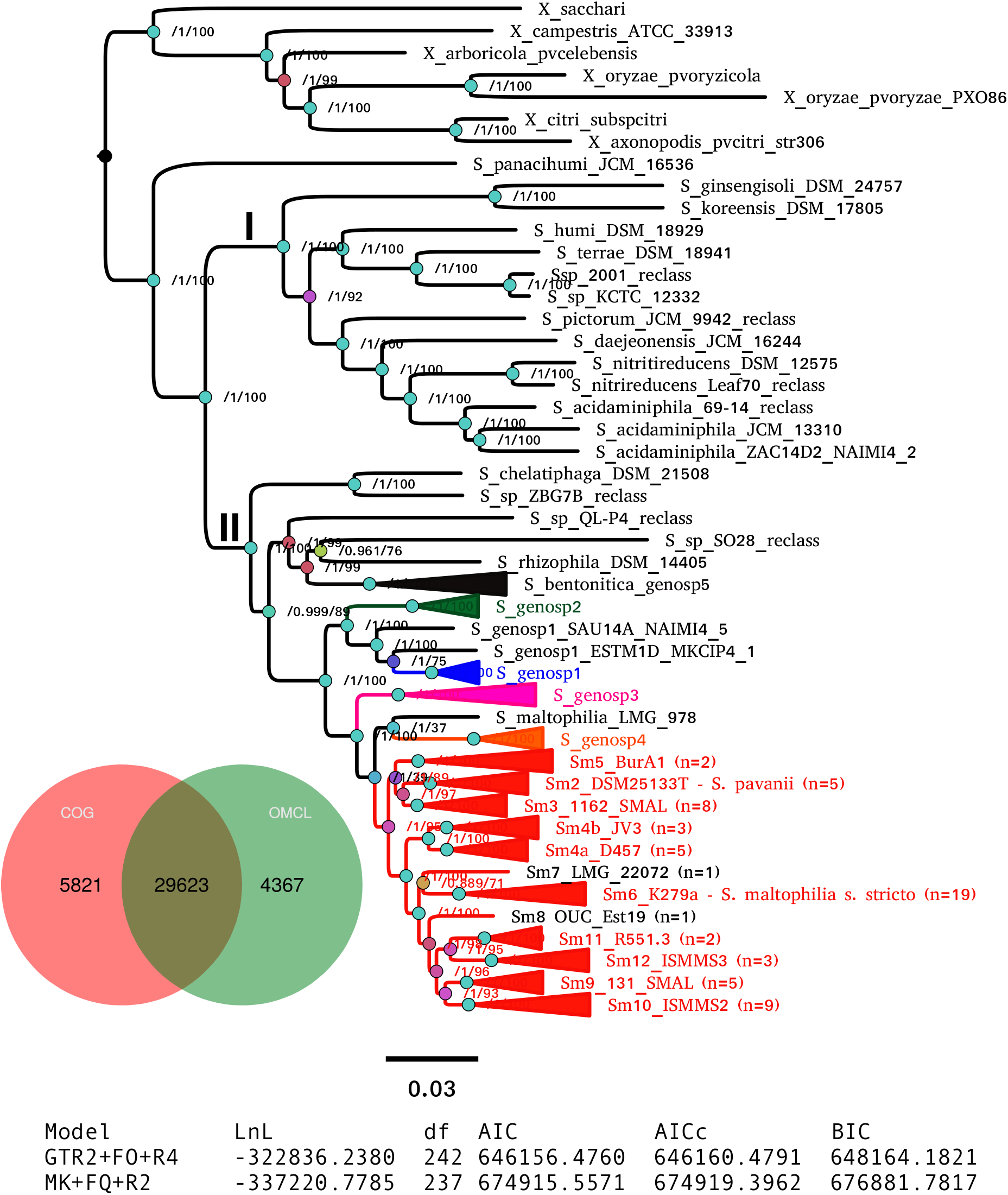
Maximum-likelihood pan-genome phylogeny estimated with IQ-TREE from the consensus pan-genome displayed in the Venn diagram. Clades of lineages belonging to the *S. maltophilia* complex are collapsed and are labeled as in **Fig. 5**. Numbers on the internal nodes represent the approximate Bayesian posterior probability/UFBoot2 bipartition support values (see methods). The tabular inset shows the results of fitting either the binary (GTR2) or morphological (MK) models implemented in IQ-TREE, indicating that the former has an overwhelmingly better fit. The scale bar represents the number of expected substitutions per site under the binary GTR2+F0+R4 substitution model.

### Application of non-supervised learning approaches to BLAST-based core-genome average nucleotide distance (cgANDb) and Gower pan-genome distances (pgGdist) provide statistically-consistent results for prokaryotic species delimitation

The final goal of any geno-taxonomic study is to identify species-like clusters. These should consist of monophyletic groups identified on genome trees that display average genome identity (gANI) values > 94 %, based on a widely accepted cutoff-value (Rosselló-Mora and Amann, 2015). In this section we searched for such species-clusters within the taxonomically problematic *Stenotrophomonas maltophilia* complex (Smc). Our core- and pan-genome phylogenies consistently identified potential species-clades within the Smc that grouped exactly the same strains (compare Figs. 5 and 6). We additionally performed a cluster analysis of core-genome ANI values computed from the pairwise BLASTN alignments (cgANIb) used to define OMCL core-genome clusters for the 86 Smc genomes analyzed in this study. The resulting cgANIb matrix was then converted to a distance matrix (cgANDb = 100 % – cgANIb) and clustered with the aid of the plot_matrix_heatmap.sh script from the GET_HOMOLOGUES suite. Figure 7 shows the resulting tree, which resolves 16 clusters within the Smc at a conservative cgANDb cutoff value of 5% (cgANIb = 95%). At this distance level, the four genospecies labeled as Sgn1-Sgn4 on Figure 5 are resolved as five clusters because the most divergent Sgn1 genome (ESTM1D_MKCIP4_1) is split as a separate lineage. This is the case also at cgANDb = 6 (Fig. 7), reason why this strain most likely represents a sixth genospecies. All these genospecies are very distantly related to the large *S. maltophilia s. lato* cluster, which gets split into 11 sub-clusters at the conservative cgANDb = 5 % cutoff. Thirteen clusters are resolved at the 4 % threshold, and a minimum of seven at the 6 % level (cgANIb = 94%), as shown by the dashed lines (Fig. 7). These results strongly suggest that the *S. maltophilia sensu lato* clade (Fig. 5) actually comprises multiple species. The challenging question is how many? In an attempt to find a statistically-sound answer, we applied an unsupervised learning approach based on the evaluation of different goodness of clustering statistics to determine the optimal number of clusters (*k*) for the cgANDb matrix. The gap-statistic and a parametric, model-based cluster analysis yielded *k* values >= 35 (data not shown). These values seem too high for this dataset, as they correspond to a gANI value > 98%. However, the more conservative average silhouette width (ASW) method (Kaufman and Rousseeuw, 1990) identified an optimal *k* = 19 (inset in Fig 7) for the complete set of Smc genomes. This number of species-like clusters is much more reasonable for this data set, as it translates to a range of cgANDb between 4.5 and 4.7 (cgANIb range: 95.5% - 95.3%). Close inspection of the ASW profile reveals that the first peak is found at *k* = 13, which has an almost identical ASW as that of the maximal value and maps to a cgANDg = 5.7 (cgANIb of 94.3%). In summary, the range of reasonable numbers of clusters proposed by the ASW statistic (*k* = 13 to *k* = 19) corresponds to cgANDb values in the range of 5.7% - 4.5% (cgANIb range: 94.3% - 95.5%), which fits well with the new gold-standard for species delimitation (gANI > 94%), established in influential works (Konstantinidis and Tiedje, 2005; Richter and Rossello-Mora, 2009). We noted however, that at a cgANDb = 4.1% (cgANIb = 95.9 %) the strain composition of the clusters was 100% concordant with the monophyletic subclades shown in the core-genome (Fig. 5) and pan-genome (Fig. 6) phylogenies. Importantly, at this cutoff, the length of the branches subtending each cluster is maximal, both on the core-genome phylogeny (Fig. 5) and on the cgANDb cladogram (Fig. 7). Based on the combined and congruent evidence provided by these complementary approaches, we can safely conclude that: *i*) the Smc genomes analyzed herein may actually comprise up to 19 or 20 different species-like lineages, and *ii*) that only the strains grouped in the cluster labeled as Sm6 in Figs. 5, 6 and 7 should be called *S. maltophilia*. The latter is the most densely sampled species-like cluster (*n* = 19) and includes ATCC 13637^T^, the type strain of the species.

**Figure 7.**
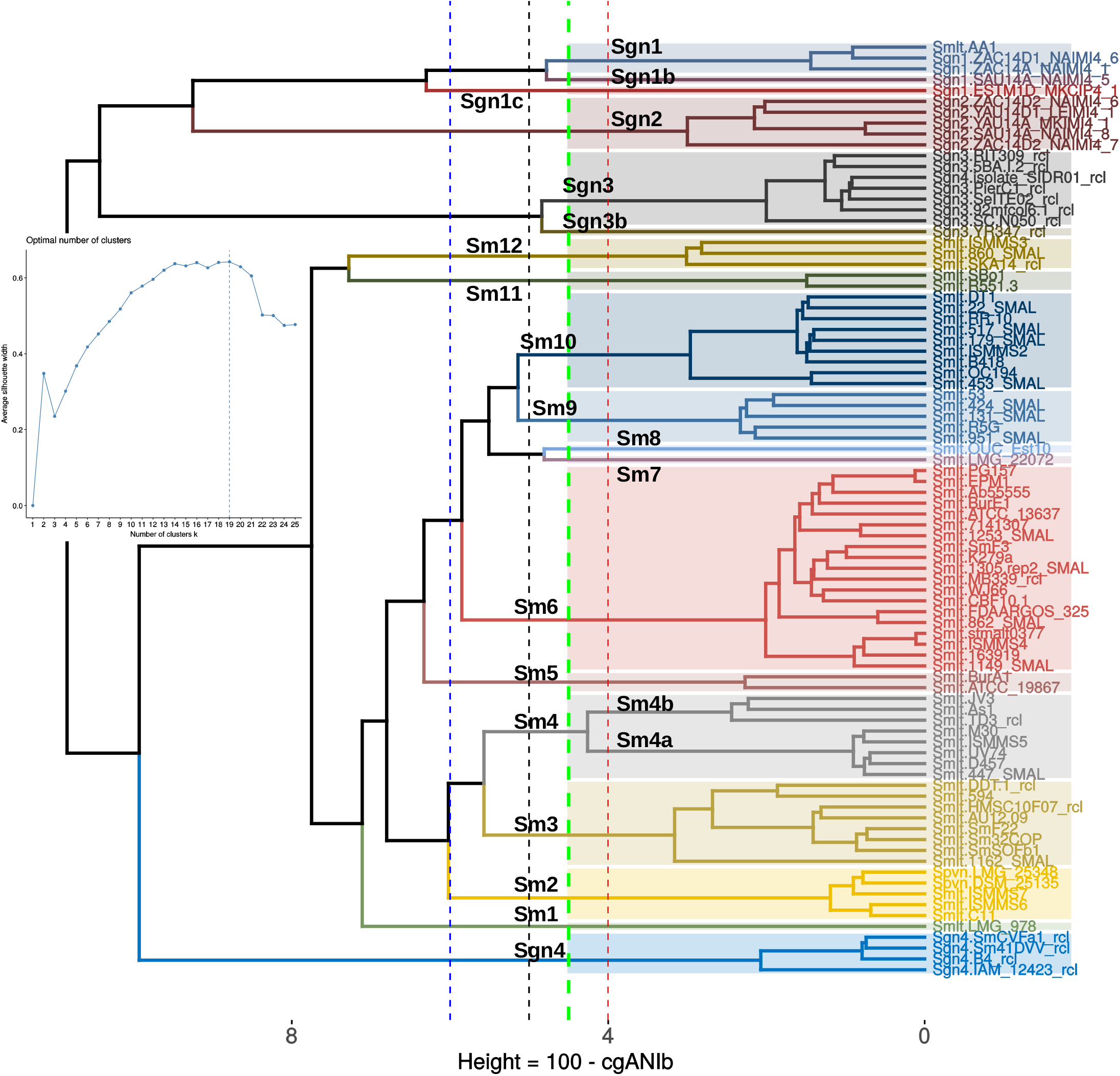
Application of an unsupervised learning approach to the cgANIb distance matrix to identify statistically-consistent species-like clusters. The cgANIb matrix was converted to a distance matrix (cgANDb) and clustered using the Ward.D2 algorithm. The optimal number of clusters (*k*) was determined with the average silhouette-width statistic. The inset shows the statistic’s profile, with *k* = 19 as the optimal number of clusters. This number corresponds to an cgANIb of 95.5 % (gray dashed line). At a cgANDb of 4.1 % (cgANIb = 95.9%) the groups delimited by the clustering approach are perfectly consistent with those delimited by the core- and pan-genome ML phylogenies displayed in **Figure 5** and **Figure 6**, respectively.

### On the ecology and other biological attributes of the species-like clusters in the *Stenotrophomonas maltophilia* complex

In this final section we present a brief summary of the ecological attributes reported for selected members of the species-like clusters resolved within the Smc (Figs 5 and 7). The four unnamed genospecies (Sgn1-Sgn4) group mainly environmental isolates. This is consistent with our previous evolutionary and ecological analyses of a comprehensive multilocus dataset of the genus (Ochoa-Sánchez and Vinuesa, 2017). In that study only Mexican environmental isolates were found to be members of the newly discovered genospecies Sgn1 and Sgn2 (named as Smc1 and Smc2, respectively). In this work we discovered that the recently sequenced maize root isolate AA1 (Niu et al., 2017), misclassified as *S. maltophilia*, clusters tightly with the Sgn1 strains (Fig. 5). The *S. maltophilia sensu lato* clade is split into 13 or 14 groups based on cgANDb (Fig 7). Sm6 forms the largest cluster, grouping mostly clinical isolates related to the type strain *S. maltophilia* ATCC 13637^T^, like the model strain K279a (Crossman et al., 2008), ISMMS4 (Pak et al., 2015), 862_SMAL, 1149_SMAL and 1253_SMAL (Roach et al., 2015), as well as EPM1 (Sassera et al., 2013), recovered from the human parasite *Giardia duodenalis.* However, this group also comprises some environmental isolates like BurE1, recovered from a bulk soil sample (Youenou et al., 2015). In summary, cluster Sm6 holds the *bona fide S. maltophilia* strains (*sensu stricto*), which may be well-adapted to associate with different eukaryotic hosts and cause opportunistic infections in humans. Cluster Sm4a contains the model strain D574 (Lira et al., 2012) along with four other clinical isolates (Conchillo-Solé et al., 2015) and therefore may represent a second clade enriched in strains with high potential to cause opportunistic pathogenic infections in humans. Noteworthy, this group is distantly related to Sm6 (Figs. 5 and 7). Cluster Sm4b is closely related to Sm4a based on the pan-genome phylogeny and the cgANDd cladogram (Figs. 6 and 7). It groups the Brazilian rhizosphere-colonizing isolate JV3, the Chinese highly metal tolerant strain TD3 (Ge and Ge, 2016) and strain As1, isolated from the Asian malaria vector *Anopheles stephensi* (Hughes et al., 2016). The lineage Sm3 holds eight isolates of contrasting origin, including the Chinese soil isolate DDT-1, capable of using DDT as the sole source of carbon and energy (Pan et al., 2016), as well as clinical isolates like 1162_SMAL (Roach et al., 2015) and AU12-09, isolated from a vascular catheter (Zhang et al., 2013), and environmental isolates like SmF22, Sm32COP and SmSOFb1, isolated from different manures in France (Bodilis et al., 2016). Cluster Sm2 groups the *S. pavanii* strains, including the type strain DSM_25135^T^, isolated from the stems of sugar cane in Brazil (Ramos et al., 2011), together with the clinical isolates ISMMS6 and ISMMS7, that carry mutations conferring quinolone resistance and causing bacteremia (Pak et al., 2015), and strain C11, recovered from pediatric cystic fibrosis patients (Ormerod et al., 2015). Cluster Sm5 includes two strains recovered from soils, ATCC 19867 which was first classified as *Pseudomonas hibiscicola*, and later reclassified as *S. maltophilia* based on MLSA studies (Vasileuskaya-Schulz et al., 2011), and the African strain BurA1, isolated from bulk soil samples collected in sorghum fields in Burkina Faso (Youenou et al., 2015). Cluster Sm9 holds clinical isolates, like 131_SMAL, 424_SMAL and 951_SMAL (Roach et al., 2015). Its sister group is Sm10. It holds 9 strains of contrasting geographic and ecological provenances, ranging from Chinese soil and plant-associated bacteria like the rice-root endophyte RR10 (Zhu et al., 2012), the grassland-soil tetracycline degrading isolate DT1 (Naas et al., 2008), and strain B418, isolated from a barley rhizosphere and displaying plant-growth promotion properties (Wu et al., 2015), to clinical isolates (22_SMAL, 179_SMAL, 453_SMAL, 517_SMAL) collected and studied in the context of a large genome sequencing project carried out at the University of Washington Medical Center (Roach et al., 2015). Cluster Sm11 tightly groups the well-characterized poplar endophyte R551-3, which is a model plant-growth-promoting bacterium (Alavi et al., 2014; Ryan et al., 2009; Taghavi et al., 2009) and SBo1, cultured from the gut of the olive fruit fly *Bactrocera oleae* (Blow et al., 2016). Cluster Sm 12 contains the environmental strain SKA14 (Adamek et al., 2014), along with the clinical isolates ISMMS3 (Pak et al., 2015) and 860_SMAL (Roach et al., 2015). Sm1, Sm7 and Sm8 each hold a single strain. The following conclusions can be drawn from this analysis: *i*) the species-like clusters within the *S. maltophilia sensu lato* (Smsl) clade (Fig. 5) are enriched in opportunistic human pathogens, when compared with the Smc clusters Sgn1-Sgn4*; ii*) most Smsl clusters also contain diverse non-clinical isolates isolated from a wide range of habitats, demonstrating the great ecological versatility found even within specific Smsl clusters like Sm3 or Sm10; *iii*) taken together, these observations strongly suggest that the Smc species-like clusters are all of environmental origin, with the potential for the opportunistic colonization of diverse human organs. This potential may be particularly high in certain lineages, like in *S. maltophilia sensu stricto* (Sm6) or Sm4a, both enriched in clinical isolates. However, a much denser sampling of genomes and associated phenotypes is required for all clusters to be able to identify statistically sound associations between them.

## DISCUSSION

In this study we developed and benchmarked GET_PHYLOMARKERS, an open-source, comprehensive, and easy-to-use software package for phylogenomics and microbial genome taxonomy. Programs like amphora (Wu and Eisen, 2008) or phylosift (Darling et al., 2014) allow users to infer genome-phylogenies from huge genomic and metagenomic datasets by scanning new sequences against a reference database of conserved protein sequences to establish the phylogenetic relationships between the query sequences and database hits. The first program searches the input data for homologues to a set of 31 highly conserved proteins used as phylogenetic markers. Phylosift is more oriented towards the phylogenetic analysis of metagenome community composition and structure. Other approaches have been developed to study large populations of a single species. These are based on the identification of single nucleotide polymorphisms in sequence reads produced by high-throughput sequencers, using either reference-based or reference-free approaches, and subjecting them to phylogenetic analysis (Timme et al., 2013). The GET_PHYLOMARKERS software suite was designed with the aim of identifying orthologous clusters with optimal attributes for phylogenomic analysis and accurate species-tree inference. It also provides tools to infer phylogenies from pan-genomes, as well as non-supervised learning approaches for the analysis of overall genome relatedness indices (OGRIs) for geno-taxonomic studies of multiple genomes. These attributes make GET_PHYLOMARKERS unique in the field.

It is well-established that the following factors strongly affect the accuracy of genomic phylogenies: *i*) correct orthology inference; *ii*) multiple sequence alignment quality; *iii*) presence of recombinant sequences; *iv*) loci producing anomalous phylogenies, which may result for example from horizontal gene transfer, differential loss of paralogues between lineages and *v*) amount of the phylogenetic signal. GET_PHYLOMARKERS aims to minimize the negative impact of potentially problematic or poorly performing orthologous clusters by explicitly considering and evaluating these factors. Orthologous clusters were identified with GET_HOMOLGOUES (Contreras-Moreira and Vinuesa, 2013) because of its distinctive capacity to compute high stringency clusters of single-copy orthologs. In this study we used a combination of BLAST alignment filtering imposing a high (90%) query coverage threshold, PFAM-domain composition scanning and calculation of a consensus core-genome from the orthologous gene families produced by three clustering algorithms (BDBH, COGtriangles and OMCL) to minimize errors in orthology inference. Multiple sequence alignments were generated with CLUSTAL-OMEGA (Sievers et al., 2012), a state-of-the-art software under constant development, capable of rapidly aligning hundreds of protein sequences with high accuracy, as reported in recent benchmark studies (Le et al., 2017; Sievers and Higgins, 2018). GET_PHYLOMARKERS generates protein alignments and uses them to compute the corresponding DNA-alignments, ensuring that the codon structure is always properly maintained. Recombinant sequences have been known for a long time to strongly distort phylogenies because they merge independent evolutionary histories into a single lineage. Recombination erodes the phylogenetic signal and misleads classic treeing algorithms, which assume a single underlying history (Didelot and Maiden, 2010; Martin, 2009; Pease and Hahn, 2013; Posada and Crandall, 2002; Schierup and Hein, 2000; Turrientes et al., 2014). Hence, the first filtering step in the pipeline is the detection of putative recombinant sequences using the very fast, sensitive and robust phi(w) statistic (Bruen et al., 2005). The genus *Stenotrophomonas* has been previously reported to have high recombination rates (Ochoa-Sánchez and Vinuesa, 2017; Yu et al., 2016). It is therefore not surprising that the phi(w) statistic detected significant evidence for recombination in up to 47% of the orthologous clusters. The non-recombinant sequences are subsequently subjected to maximum-likelihood phylogenetic inference to identify anomalous trees using the non-parametric *kdetrees* statistic (Weyenberg et al., 2014, 2017). The method estimates distributions of phylogenetic trees over the "tree space" expected under the multispecies-coalescent, identifying outlier trees based on their topologies and branch lengths in the context of this distribution. Since this test is applied downstream of the recombination analysis, only a modest, although still significant proportion (14%-17%) of outlier trees were detected (Table 2). The next step determines the phylogenetic signal content of each gene tree (Vinuesa et al., 2008). It has been previously established that highly informative trees are less prone to get stuck in local optima (Money and Whelan, 2012). They are also required to properly infer divergence at the deeper nodes of a phylogeny (Salichos and Rokas, 2013), and to get reliable estimates of tree congruence and branch support in large concatenated datasets typically used in phylogenomics (Shen et al., 2017). We found that IQ-TREE-based searches allowed a significantly more efficient filtering of poorly resolved trees than FastTree. This is likely due to the fact that the former fits more sophisticated models (with more parameters) to better account for among-site rate variation. Under-parameterized and poorly fitting substitution models partly explain the apparent overestimation of bipartition support values done by FastTree. This is also the cause of the poorer performance of FastTree, which finds gene trees that generally have lower ln*L* scores than those found by IQ-TREE. A recent comparison of the performance of four fast ML phylogenetic programs using large phylogenomic data sets identified IQ-TREE (Nguyen et al., 2015) as the most accurate algorithm. It consistently found the highest-scoring trees. FastTree (Price et al., 2010) was, by large, the fastest program evaluated, although at the price of being the less accurate one (Zhou et al., 2017). This is in line with our findings. We could show that the higher accuracy of IQ-TREE is particularly striking when using large concatenated datasets. As stated above, this is largely attributable to the much richer choice of models implemented in the former. ModelFinder (Kalyaanamoorthy et al., 2017) selected GTR+ASC+F+R6 model for the concatenated supermatrix, which is much richer in parameters than the GTR+CAT+Gamma20 model fitted by FastTree. The +ASC is an ascertainment bias correction parameter, which should be applied to alignments without constant sites (Lewis, 2001), such as the supermatrices generated by GET_PHYLOMARKERS (see methods). The FreeRate model (+R) generalizes the +G model (fitting a discrete Gamma distribution to model among-site rate variation) by relaxing the assumption of Gamma-distributed rates (Yang, 1995). The FreeRate model typically fits data better than the +G model and is recommended for the analysis of large data sets (Soubrier et al., 2012).

The impact of substitution models in phylogenetics has been extensively studied (Posada and Buckley, 2004). However, the better models implemented in IQ-TREE are not the only reason for its superior performance. A key aspect strongly impacting the quality of phylogenomic inference with large datasets is tree-searching. This has been largely neglected in most molecular systematic and phylogenetic studies of prokaryotes (Ochoa-Sánchez and Vinuesa, 2017; Vinuesa, 2010; Vinuesa et al., 2008). Due to the factorial increase of the number of distinct bifurcating topologies possible with every new sequence added to an alignment (Felsenstein, 2004a), searching the tree-space for large datasets is an NP-hard (non-deterministic polynomial-time) problem that necessarily requires heuristic algorithms. This implies that once an optimum is found, there is no way of telling whether it is the global one. The strategy to gain quantitative evidence about the quality of a certain tree is to compare its score in the context of other trees found in searches initiated from a pool of different seed trees. Due to the high dimensionality of the likelihood space, and the strict “hill-climbing” nature of ML tree search algorithms (Felsenstein, 2004a), they generally get stuck in local optima (Money and Whelan, 2012). The scores of the best trees found in each search can then be compared in the form of an “ln*L* score profile”, as performed in our study. Available software implementations for fast ML tree searching use different branch-swapping strategies to try to escape from early encountered “local optima”. IQ-TREE implements a more efficient tree-searching strategy than FastTree, based on a combination of hill-climbing and stochastic nearest-neighbor interchange (NNI) operations, always keeping a pool of seed trees, which help to escape local optima (Nguyen et al., 2015). This was evident when the ln*L* score profiles of both programs were compared. IQ-TREE found a much better scoring species tree despite the much higher number of independent searches performed with FastTree (50 vs. 1001) using its most intensive branch-swapping regime. An important finding of our study is the demonstration that the ln*L* search profile of IQ-TREE is much shallower than that of FastTree. This suggests that the former finds trees much closer to the potential optimum than the latter. It has been shown that the highest-scoring (best) trees tend to have shorter branches, and overall tree-length, than those stuck in worse local optima (Money and Whelan, 2012). In agreement with this report, the best species-tree found by IQ-TREE has a notoriously shorter total length and significantly shorter edges than those of the best species-tree found by FastTree.

Our extensive benchmark analysis conclusively demonstrated the superior performance of IQ-TREE. Based on this evidence, it was chosen as the default search algorithm for GET_PHYLOMARKERS. However, it should be noted that topological differences between the best trees found by both programs were minor, not affecting the composition of the major clades in the corresponding species trees. It is therefore safe to conclude that the reclassification of *Stenotrophomonas* genome sequences proposed in Table 3 is robust. They are consistently supported by the species-trees estimated with both programs. This result underlines the utility of GET_PHYLOMARKERS to identify misclassified genomes in public sequence repositories, a problem found in many genera (Gomila et al., 2017; Sangal et al., 2016). GET_PHYLOMARKERS is unique in its ability to combine core-genome phylogenomics with ML and parsimony phylogeny estimation from the pan-genome matrix. In line with other recent studies (Caputo et al., 2015; Tu and Lin, 2016), we demonstrate that pan-genome analyses are valuable in the context of microbial molecular systematics and taxonomy. All genomes found to be misclassified based on the phylogenomic analysis of core-genomes were corroborated by the ML and parsimony analyses of the PGM. Furthermore, the combined evidence gained from these independent approaches consistently revealed that the Smc contains up to 20 monophyletic and strongly supported species-like clusters. These are defined at the cgANIb 95.9% threshold, and include the previously identified genospecies Smc1-Smc4 (Ochoa-Sánchez and Vinuesa, 2017), and up to 13 genospecies within the *S. maltophilia sensu lato* clade. This threshold fits well with the currently favored gANI > 94% cutoff for species delimitation (Konstantinidis and Tiedje, 2005; Richter and Rossello-Mora, 2009). The consistency among all the different approaches strongly supports the proposed delimitations. We used an unsupervised learning procedure to determine the optimal number of clusters (*k*) in the cgANDb matrix computed from the 86 Smc genomes analyzed. The average silhouette width goodness of clustering statistic proposed an optimal *k* = 19, which corresponds to a gANI = 95.5%. At this cutoff, 13 (instead of 14) species-like clusters are delimited within the *S. maltophilia sensu lato* clade. This unsupervised learning method therefore seems promising to define the optimal number of clusters in ANI-like matrices using a statistical procedure. However, it should be critically and extensively evaluated in other geno-taxonomic studies to better understand its properties and possible limitations, before being broadly used.

Current models of microbial speciation predict that bacterial species-like lineages should be identifiable by significantly reduced gene flow between them, even when recombination levels are high within species (Cadillo-Quiroz et al., 2012; Shapiro et al., 2012). Such lineages should also display differentiated ecological niches and phenotypes (Koeppel et al., 2008; Shapiro and Polz, 2015). In our previous comprehensive multilocus sequence analysis of species borders in the genus *Stenotrophomonas* (Ochoa-Sánchez and Vinuesa, 2017) we could show that those models fitted our data well. We found highly significant genetic differentiation and marginal gene-flow across strains from sympatric Smc1 and Smc2 lineages, as well as highly significant differences in the resistance profiles of *S. maltophilia sensu lato* isolates versus Smc1 and Smc2 isolates. We could also show that all three lineages have different habitat preferences (Ochoa-Sánchez and Vinuesa, 2017). The genomic analyses presented in this study for five Smc1 and Smc2 strains, respectively, fully support their separate species status from a geno-taxonomic perspective. Given the recognized importance of gene gain and loss processes in bacterial speciation and ecological specialization (Caputo et al., 2015; Jeukens et al., 2017; Richards et al., 2014; Shapiro and Polz, 2015), as reported also in plants (Gordon et al., 2017), we think that the evidence gained from pan-genome phylogenies is particularly informative for microbial geno-taxonomic investigations. We believe they should be used to validate the groupings obtained by the classical gANI cutoff-based species delimitation procedure (Goris et al., 2007; Konstantinidis and Tiedje, 2005; Richter and Rossello-Mora, 2009) that dominates current geno-taxonomic research. It is well documented that pan-genome-based groupings tend to better reflect ecologically relevant phenotypic differences between groups (Caputo et al., 2015; Jeukens et al., 2017; Lukjancenko et al., 2010). We recommend that future geno-taxonomic studies search for a consensus of the complementary views of genomic diversity provided by OGRIs, core- and pan-genome phylogenies, as performed herein. GET_PHYLOMARKERS is a useful and versatile tool for this task.

In summary, in this study we developed a comprehensive and powerful suite of open-source computational tools for state-of-the art phylogenomic and pan-genomic analyses. Their application to critically analyze the geno-taxonomic status of the genus *Stenotrophomonas* provided compelling evidence that the taxonomically ill-defined *S. maltophilia* complex holds many cryptic species. However, we refrain at this point from making formal taxonomic proposals for them because we have not yet performed the above-mentioned population genetic analyses to demonstrate the genetic cohesiveness of the individual species and their differentiation from closely related ones. This will be the topic of a follow-up work in preparation. We think that comparative genomic analyses designed to identify lineage-specific genetic differences that may underlie niche-differentiation of species are the most powerful and objective criteria to delimit species in any taxonomic group (Ochoa-Sánchez and Vinuesa, 2017; Vinuesa et al., 2005).

## AUTHOR CONTRIBUTIONS

PV designed the project, wrote the bulk of the code, assembled the genomes, performed the analyses and wrote the paper. LEOS isolated the strains sequenced in this study and performed all wet-lab experiments. BCM was involved in the original design of the project, contributed code, and set up the docker image. All authors read and approved the final version of the manuscript.

## ACKNOWLEDGEMENTS

We thank Javier Rivera for excellent technical support with wet-lab experiments and José Alfredo Hernández and Víctor del Moral for support with server administration. Jason Steeel from the DNASU Sequencing Core at The Biodesign Institute, Arizona State University, is acknowledged for generating the genome sequences of our samples. Dr. Claudia Silva is thanked for her critical reading of the manuscript. We are thankful to GitHub (https://github.com/), docker (https://hub.docker.com/) and the open-source community at large, for providing great resources for software development.

